# Harnessing distinct tissue-resident immune niches *via* S100A9/TLR4 improves ketone, lipid, and glucose metabolism

**DOI:** 10.1101/2025.05.21.655283

**Authors:** Giulia Lucibello, Gloria Ursino, Pryscila D. S. Teixeira, Szabolcs Zahoran, Francesca Fanuele, Marinos Kallikourdis, Florian Visentin, Christelle Veyrat-Durebex, Ariane Widmer, Yibo Wu, Marco Cremonesi, Claes B. Wollheim, Perrine Castets, Giorgio Ramadori, Roberto Coppari

**Author notes:** These authors contributed equally to this work (Giulia Lucibello, Gloria Ursino).

## Abstract

Immunometabolism contributes to the development of metabolic diseases. Yet, how certain metabolic disorders, such as insulin deficiency (ID), influence immune cell function is poorly understood. Here, we observe that ID rearranges the immune landscape of the liver, causing a decrease in T cells and an increase in Kupffer cells, accompanied by a shift in the transcriptome and polarization of the latter. Treating ID mice with the protein S100A9 rescues the polarization and lipid-related changes caused by ID in the KCs, and rescues hypertriglyceridemia and hyperketonemia in a TLR4-dependent manner. Additionally, S100A9 acts on other immune niches to increase glucose uptake in skeletal muscle, improving hyperglycemia. In summary, the S100A9-TLR4 axis is a new tool to harness immune cells for improving ID-related metabolic dysfunction.

## Introduction

Immunometabolism is an emerging field focusing on the interplay between immunological and metabolic pathways^1^. In the past three decades, it has become increasingly evident that immune cells exert a key role in maintaining metabolic homeostasis. When their activity is altered, they can contribute to the development of metabolic diseases^2,3^. For example, the physiological secretion of insulin from pancreatic beta (β)-cells is regulated by the plasma glucose content and several other stimuli, including myeloid-cell-derived IL-1β^4,5^. However, type 2 diabetes mellitus (T2DM) patients display macrophage infiltration of the endocrine pancreas^6^ and chronic elevation of IL-1β contributes to β-cells damage^6^. Also, increased production of inflammatory cytokines contributes to the decreased insulin secretion occurring before the onset of type 1 diabetes mellitus (T1DM)^7^. In obesity and T2DM, macrophages and dendritic cells accumulate in adipose tissue^8^. In this context, these cells acquire a pro-inflammatory profile and secrete increased amounts of inflammatory cues (e.g., tumor necrosis factor alpha, TNFα), contributing to the low-grade inflammatory state that is characteristic of these illnesses^9^. Moreover, in the hypothalamus, specific neurons are empowered with metabolic-sensing mechanisms (e.g., their activity is influenced by the amount of circulating cues as for example insulin, leptin and other hormones) and are key for metabolic homeostasis^10^. The function of these neurons is influenced by hypothalamic macrophages, whose profile is shaped by the nutritional input. In obesity, their pro-inflammatory profile impairs the ability of hypothalamic neurons to properly respond to peripheral signals and hence contributes to the development of metabolic imbalance^11^. In a healthy liver, resident macrophages (also known as Kupffer cells; KCs represent about 15% of the hepatic cell mass) are located intravascularly and line the fenestrated sinusoidal vessels^12^. KCs have self-renewal and phagocytic capacity and are well known for their detoxifying functions (e.g., they remove damaged cells and engulf pathogens and other toxic products of digestion)^13^. In addition to these functions, KCs are also involved in physiological metabolic programs, as they contribute to fasting-induced ketogenesis *via* reduced TNFα secretion^14^. KCs are heterogeneous and at least two different subsets have been reported^15,16^. The KC1 subset is characterized by the low expression of CD206 and does not express Endothelial Cell Adhesion Molecule (ESAM), while the KC2 subset displays high CD206/CD36 level and expresses ESAM ^15,16^. Of note, upon fat-rich feeding, the KC2 subset becomes more abundant and active, and contributes to the development of fat-rich-diet-induced obesity and hepatic steatosis^15^. Skeletal muscle (SkM) also possesses a rich resident immune environment^17^. These cells are known to play important roles in regulating SkM growth, regeneration, and repair in injury or disease contexts^18–20^, and in response to physiological inputs such as exercise^21,22^. Yet the modality by which they regulate metabolic processes (e.g., glucose and lipid metabolism) in myocytes is unknown. Noteworthy, SkM of obese and insulin-resistant patients feature increased resident and locally recruited T-cells and macrophages, which tend to polarize towards pro-inflammatory phenotypes thus contributing to a low-grade inflammatory state^23,24^.

Although tissue-resident immune niches are involved in the pathophysiology of metabolic diseases (e.g., obesity, diabetes, hepatic steatosis), means to harness them for therapeutic purposes are lacking. For example, earlier studies suggesting a crucial role of TNFα have spurred several TNFα-blockade clinical studies in individuals with metabolic syndrome or diabetes^25,26^. However, they failed to improve insulin sensitivity and glucose metabolism^27–29^. Moreover, although clinical trials with IL-1β antagonists showed beneficial effects on cardiovascular events in patients with prior myocardial infarction, they failed to improve glycemic control in diabetic subjects or to prevent new onset diabetes^29^.

Here, we focused on a metabolic condition affecting millions worldwide; namely insulin deficiency (ID)^30,31^. ID is a heterogeneous condition that develops following surgical removal of the pancreas, autoimmune-mediated attack of pancreatic β-cells (as in T1DM) or pancreatic β-cell exhaustion, dedifferentiation, and/or death due to metabolic stress (as in T2DM)^31^. While its etiology might differ, ID is characterized by several metabolic derangements, including but not limited to hyperglycemia, dyslipidemia, and hyperketonemia. If untreated, ID is a lethal condition. Its treatment is based on insulin therapy, which, however, is suboptimal as it exposes the patients to life-threatening hypoglycemia and does not protect ID patients from having a higher risk of severe complications, as for example cardiovascular disease, retinopathy, and nephrophaty^32^.

By generating and assessing diverse genetically engineered animal models of ID, we discovered a previously unknown role for the KCs and bone marrow (BM) derived immune cells in mediating the beneficial action of the Ca^2+^-binding protein S100A9 on lipid and ketone metabolism in the liver, and on glucose metabolism in SkM, respectively. By affecting different immune niches in the liver and SkM S100A9 rescues hypertriglyceridemia and hyperketonemia and stimulates SkM glucose uptake *via* Toll-like-receptor 4 (TLR4)-dependent mechanism. These results indicate that targeting distinct immune cells by the S100A9-TLR4 pathway improves the key metabolic imbalances caused by ID.

## Results

### Insulin deficiency reshapes the immunophenotype of the liver

Despite the key role of immunometabolism in the liver^13–15^ and insulin’s central role in hepatic metabolic regulation^33–35^, the consequences of ID on the hepatic immune landscape are unknown. To model human ID in mice, we used an established method^36–39^. Briefly, we employed the *RIPDTR* allele that carries the *Diphtheria Toxin receptor* (*DTR*) sequence downstream of the *rat insulin promoter* (*RIP*), allowing pancreatic β-cell depletion and consequential ID upon Diphtheria toxin (DT) administration^8^. First, we assessed whether ID affects the composition of the hepatic immune cell population. To this end, we performed fluorescence-activated cell sorting (FACS) analysis of liver samples obtained from ID mice (i.e., DT-treated *RIPDTR* mice) and healthy controls (i.e., gender-and age-matched DT-untreated *RIPDTR* mice) (Fig. 1A). In line with DT-induced pancreatic β-cell depletion^8^, ID mice reached undetectable levels of circulating insulin (Fig. S1A).

**Fig. 1.**
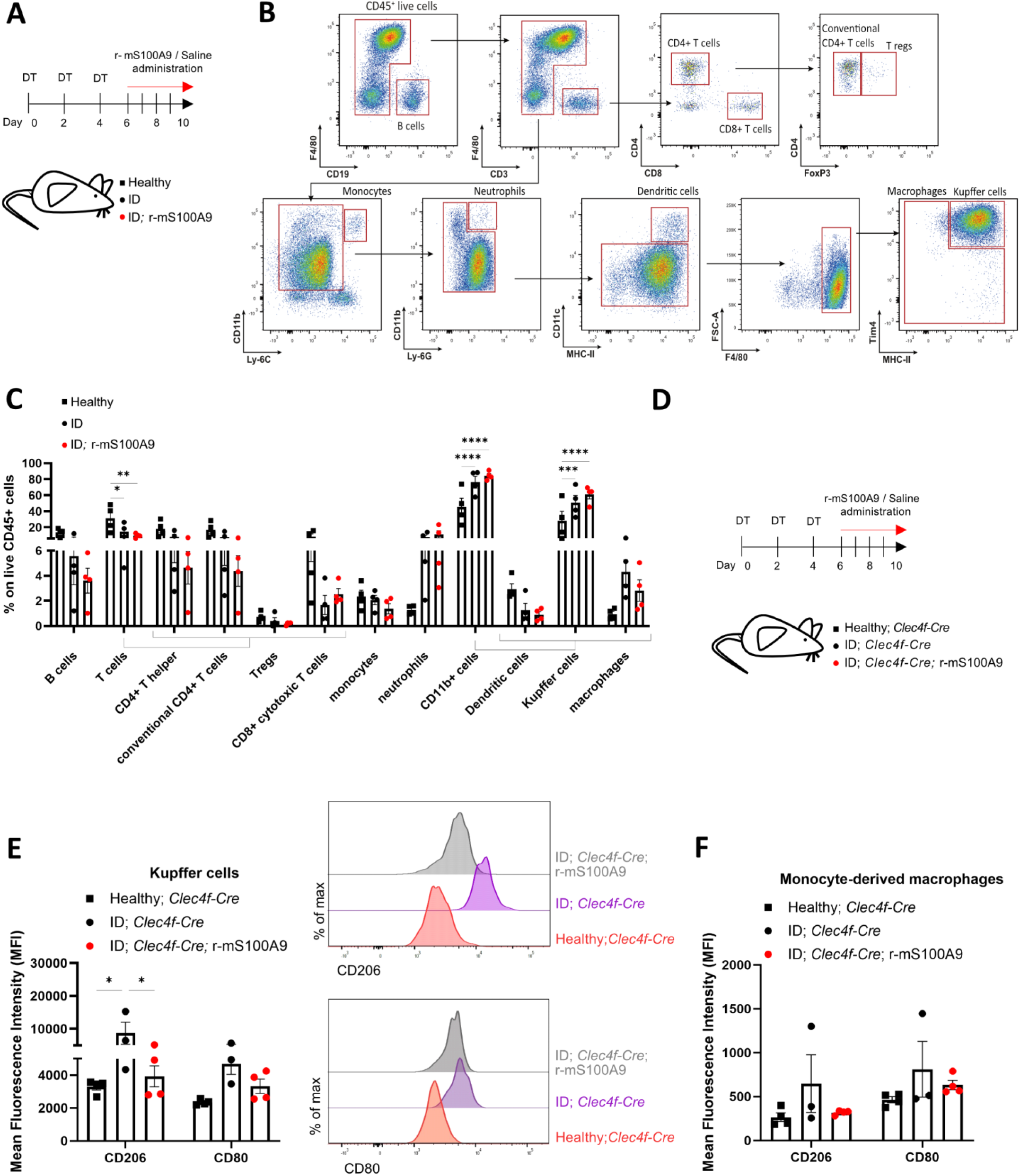
Insulin deficiency reshapes the immunophenotype of the liver. **(A)** Schematic representation of experimental groups and treatment: Healthy (n=4), ID (n=4), and ID; r-mS100A9 (n=4). To reach ID, age-and gender-matched *RIPDTR* mice received three DT injections on days 0, 2, and 4. Starting from day 6, ID groups were administered 2 times/day with either r-mS100A9 (ID; r-mS100A9) or saline (ID). On day 10, mice were sacrificed and hepatic cells were collected after hepatic perfusion. **(B)** Flow cytometry plots showing the gating strategy used to identify the immune populations of the liver (CD19+ B cells, CD4+ T cells, CD8+ T cells, FoxP3+ Tregs, CD11b+ Ly-6C+ monocytes, CD11b+ Ly-6G+ neutrophils, CD11c+ dendritic cells, Tim4+ MHCII+ Kupffer cells, F4/80+ macrophages). **(C)** Immunophenotyping of the liver in the groups as in (**A**). Percentages represent the percentage of the indicated cell type over the total amount of living CD45+ cells. **(D)** Age-and gender-matched *Clec4f-Cre* mice were rendered ID through 3 DT injections (days 0, 2, and 4) and administered with saline (ID; *Clec4f-Cre*, n=3) or r-mS100A9 (ID; *Clec4f-Cre*; r-mS100A9, n=4) for 4 days (days 6 to 10). On day 10, ID; *Clec4f-Cre* and ID; *Clec4f-Cre*; r-mS100A9 mice and their control Healthy*; Clec4f-Cre* mice (n=4) were sacrificed, and after hepatic perfusion, hepatic immune cells were collected for cytofluorimetric analysis. **(E)** Mean Fluorescence Intensity (MFI) in the KC population with flow cytometry profiles of the intensity of expression of CD206 (right, top) and CD80 (right, bottom) of mice as in (**D**). **(F)** Mean Fluorescence Intensity (MFI) in the Monocyte-derived macrophages for the indicated markers (CD206, CD80) of mice as in (**D**). Error bars represent SEM, statistical analyses were done using one-way or two-way ANOVA (Tukey’s post-hoc test). Comparisons were made with Healthy controls unless otherwise specified on the graph. *p ≤ 0.05, **p ≤ 0.01, ***p ≤ 0.001, ****p≤0.0001.

Overall, ID induced a reduction in all the immune non-macrophage cell types in the liver (identified as previously done^40^). In particular, there was a significant decrease in the T cells compartment [including CD4+ T cells, CD8+ T cells, and FoxP3+ regulatory T cells (T regs)], in favor of a significant increase in the CD11b+ positive fraction (which includes CD11c+ dendritic cells, and F4/80+ macrophages). Specifically, there was a marked increase in the liver-resident macrophages, the KCs, identified as Tim4^+^ MHC-II^+^ cells ^40^ (Figs. 1B-C). Recent studies identified more than one KC subset with different polarization profiles and different functions^12,15,16^. Thus, we set to determine the impact of ID on the polarization profile of the KCs. To this end, we crossed the *RIPDTR* allele with the *Clec4f-Cre-tdTomato* allele, a knock-in allele in which the C-type lectin domain family 4 member f, *Clec4f*, promoter drives expression of Cre-recombinase and tdTomato only in KCs^41^. Taking advantage of the KC-restricted TdTomato expression, we used FACS to measure the level of expression of selected macrophage markers in the KC population (defined by the canonical macrophage markers F4/80^+^ and CD11b^+40^, plus TdTomato^+^, Fig. S1B) and monocyte-derived macrophages (MoMΦ, defined as previously done as F4/80^+^ CD11b^+40^, and TdTomato^-^, Fig. S1B) in the liver of ID (DT-treated *RIPDTR* mice also bearing the *Clec4f-Cre-tdTomato* allele, referred here as ID; *Clec4f-Cre*) and healthy (i.e., gender-and age-matched DT-untreated *RIPDTR* mice also bearing the *Clec4f-Cre-tdTomato* allele, referred here as healthy; *Clec4f-Cre*) controls (Fig. 1D, S1B). In keeping with DT-induced pancreatic β-cell depletion^8^, ID; *Clec4f-Cre* mice displayed undetectable levels of circulating insulin compared to their controls (Fig. S1C). KCs isolated from ID*; Clec4f-Cre* mice had increased levels of the polarization marker CD206 and a trend of increased CD80 compared to their healthy controls (Fig. 1E). CD206 level on the KCs has been used to differentiate between KC subsets deputed to different functions^15,16^. In particular, CD206^HIGH^ KCs (referred to as KC2 subset) have been related to lipid metabolism in the liver, and KC2 are recruited in other lipid-rich contexts such as diet-induced obesity^15^. Whereas, CD206^LOW^ KCs (referred to as KC1 subset) display more of an immune signature^15^. CD80 is a costimulatory molecule, mostly linked with the antigen-presentation capacity of the macrophages, whose expression was previously found to be increased in the same CD206^HIGH^ KC population^15,16^. On the other hand, MoMΦ isolated from ID*; Clec4f-Cre* mice had a similar level of CD206 and CD80 compared to their healthy controls (Fig. 1F). Taken together, these results indicate that ID rearranges the immune cell populations in the liver, where T cell presence lowers while the KC component increases, accompanied by a shift of polarization of the latter towards the KC2 phenotype.

### Unbiased assays unmask KC transcriptomic changes caused by insulin deficiency

To delve deeper into the changes occurring in the KCs in ID, we performed an unbiased transcriptional characterization of the KCs. To perform RNA-sequencing (RNA-seq) analysis of KCs specifically, we crossed the “isolation of nuclei tagged in specific cell types” (INTACT) allele (*CAG-Sun1/sfGFP* mice^42^) with the *RIPDTR; Clec4f-Cre* mice. Mice bearing the *CAG-Sun1/sfGFP*, *Clec4f-Cre-tdTomato,* and *RIPDTR* alleles (referred herein as KC-INTACT mice) were generated (Fig. S2A). Immunohistochemical data shown in Fig. S2B indicate that 40% of cells expressing F4/80 (an established marker of macrophages^12^) express the Green Fluorescent Protein (GFP) in their nuclei in the liver of KC-INTACT mice. Importantly, virtually 100% of the cells expressing GFP were positive for F4/80 in the liver of KC-INTACT mice (Fig. S2B), hence indicating eutopic expression of GFP in this model. To further validate this animal model, KCs and hepatocytes were isolated from the liver of *Clec4f-Cre* mice (*RIPDTR; Clec4f-Cre* without the *Sun1/sfGFP* allele; hepatocytes and KCs from these mice are referred to as Hep^WT^ and KC^WT^, respectively) and KC-INTACT mice (hepatocytes and KCs from these mice are referred to as Hep^GFP^ and KC^GFP^, respectively) (see methods section for details). As shown in Fig. S2C, *Gfp* mRNA content is highly enriched in KC^GFP^. Collectively, these data establish that KC-INTACT mice express GFP in the nuclei of KCs specifically and thus validate our animal model.

KC-INTACT mice were rendered ID through DT treatment (ID; KC-INTACT) (Fig. 2A). In line with the expected pancreatic β-cell depletion, ID; KC-INTACT mice displayed undetectable levels of circulating insulin compared to their healthy (i.e., gender-and age-matched, DT-untreated KC-INTACT mice) controls (Fig. S2D). KC nuclei were isolated (as previously described^21^) and utilized for RNA-seq analysis. This assay identified 298 genes whose expression was significantly differentially expressed in ID; KC-INTACT mice compared to their healthy; KC-INTACT controls, of which 129 were upregulated (log_2_ fold change>1.2) and 169 downregulated (log_2_ fold change <-1.2) (Fig. S2E, Figs. 2B-D). Among the genes downregulated in ID, there was functional enrichment for genes involved in antigen processing and presentation (e.g. *Cd74, H2-Aa, H2-Ab1, H2-Eb1*), unsaturated fatty acid metabolism (e.g. *Cd74, Cyp4a12a, Elovl3, Scd1*), and regulation of glucose metabolism (e.g. *Gck, Mup16, Mup3, Serpina12*) (Fig. 2C, left). Interestingly, performing functional enrichment in the Reactome database^43^, among the most significant terms we found PPARγ and RXRα binding their target gene loci (e.g. *H2ac8, H2bc14, H2bc8, H3c13, H3c3, H4c2, Thrsp, Scd1*) (Fig. 2C, right). Both PPARγ and RXRα are transcription factors associated with the regulation of immune response, lipid, and glucose metabolism^44–46^. On the other hand, the genes that were identified as significantly upregulated in ID are involved in amino acid metabolism (e.g. *Aass, Acot4, Adh7, Agxt, Aldh1l1, Ehhadh, Gldc, Got1, Hgd, Hpd, Irs2, Kyat3, Plin5, Sesn2, Uroc1*), fatty acid oxidation and lipid modification (e.g. *Adh7, Dgka, Ehhadh, Fmo1, Fmo2, Irs2, Pdk4, Plin5, Sesn2*) (Fig. 2D, left). Whereas, when observing the Reactome in which these genes are involved, the most enriched terms involved mitochondrial metabolism (CLPXP and LONP1 binding to mitochondrial matrix proteins, e.g. *Acot1, Acot3, Fh1*) (Fig. 2D, right). This is in line with previously published data indicating that polarization changes in macrophages are accompanied by a switch between glycolytic and oxidative metabolism, therefore suggesting different mitochondrial activity^47^. Collectively, our data demonstrate that the transcriptional signature of KCs is reshaped by ID.

**Fig. 2.**
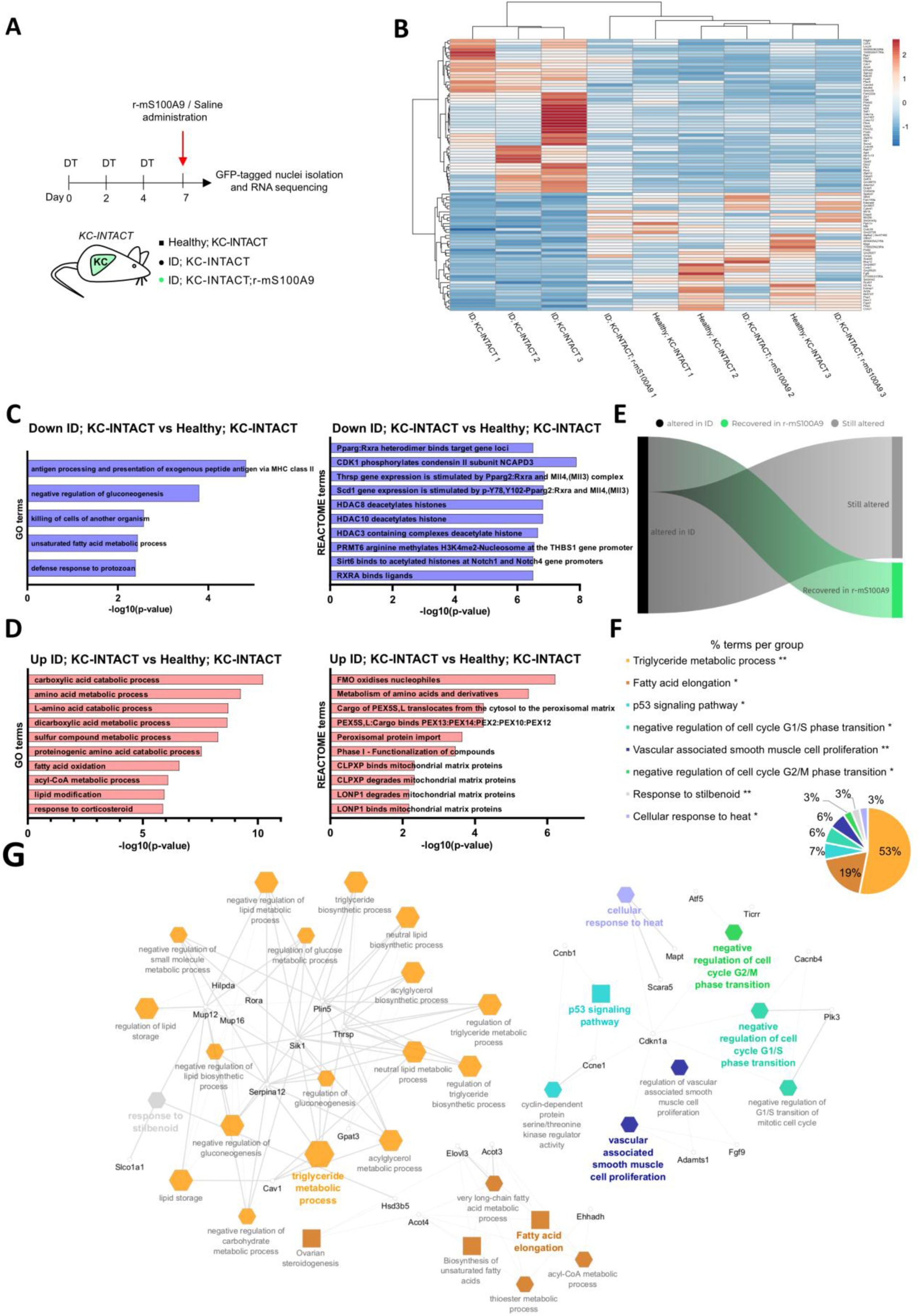
S100A9 rescues KC polarization and lipid-related transcriptional alterations caused by ID. **(A)** Scheme indicating the generation and treatment of the experimental groups: age-and gender-matched Healthy; KC-INTACT (n=3), ID; KC-INTACT (n=3), and ID; KC-INTACT; r-mS100A9 (n=3). KC-INTACT mice received three DT injections on days 0, 2, and 4 to reach ID. On day 7 they were administered with saline (ID; KC-INTACT) or r-mS100A9 (ID; KC-INTACT; r-mS100A9). 3h after saline or r-mS100A9 administration, the GFP-tagged nuclei were isolated and analyzed by RNA-seq. **(B)** Heatmap showing the normalized counts for significant (p<0.05) transcripts differentially expressed in ID; KC-INTACT versus Healthy; KC-INTACT (log2 fold change<-2 or >2) and not differentially expressed in ID; KC-INTACT; r-mS100A9 versus Healthy; KC-INTACT (-2< log2 fold change<2). Created using Heatmapper.ca **(C**-**D)** ClueGO functional enrichment analysis of the transcripts downregulated (**C**) or upregulated (**D**) in ID; KC-INTACT compared to Healthy; KC-INTACT according to the GO terms and REACTOME databases. **(E)** Infogram about the proportion of genes altered in the ID; KC-INTACT and rescued or not in the ID; KC-INTACT; r-mS100A9 group. **(F)** Pie chart showing ClueGO functional enrichment analysis of the transcripts reported in the heatmap shown in (**B**). Percentages represent the percentage of transcripts in each functional group. **(G)** Network showing ClueGO functional enrichment analysis of the transcripts reported in the heatmap shown in (**B**). Hexagons represent selected GO Biological processes, ellipses represent GO Immune system process, and rectangles represent KEGG pathways. Edge thickness is based on Kappa score and node size is based on term p-value with Bonferroni correction. Genes from highlighted GO and KEGG terms are represented in black.

### S100A9 rescues KC polarization and lipid-related transcriptional alterations caused by ID

We previously identified the Ca^2+^-binding protein S100A9 as an anti-diabetic agent able to improve hyperglycemia (by stimulating glucose uptake in SkM^39^), hyperketonemia [by suppressing hepatic fatty acid oxidation *via* Toll-Like Receptor 4 (TLR4) expressed in hepatic non-parenchymal cells^38^], and hypertriglyceridemia (*via* an unknown mechanism) caused by ID. Given that S100A9 is an immunomodulatory protein^48,49^, we directly tested whether recombinant murine S100A9 (r-mS100A9) administration to ID mice affects the hepatic immune landscape. To this end, r-mS100A9 was administrated to ID mice (i.e., DT-treated *RIPDTR* mice that underwent r-mS100A9 administration referred to as ID; r-mS100A9 mice, Fig. 1A), as previously done^38,39^. Similar to the ID group, ID; r-mS100A9 mice reached undetectable levels of circulating insulin (Fig. S1A). FACS analysis revealed a similar decrease in immune non-macrophage cell types (particularly the T cell compartment) accompanied by a similar increase in the CD11b+ positive fraction and KCs in ID; r-mS100A9 and ID mice as compared to their healthy controls (Figs. 1B, C). To better determine the effect of r-mS100A9 on KCs, DT-treated *RIPDTR* mice also bearing the *Clec4f-Cre-tdTomato* allele that underwent r-mS100A9 administration (denoted as ID; *Clec4f-Cre*; r-mS100A9 mice) were generated (Fig. 1D). Similar to the ID; *Clec4f-Cre* group, ID; *Clec4f-Cre*; r-mS100A9 mice reached undetectable levels of circulating insulin (Fig. S1C). We then compared expression levels of the polarization markers altered in ID (CD206 and CD80) between the ID; *Clec4f-Cre*; r-mS100A9 group and ID; *Clec4f-Cre* mice and their Healthy; *Clec4f-Cre* controls (Fig. 1E, Fig. S1B). Of note, data shown in Fig. 1E demonstrate that the expression levels of CD206 and CD80 are increased in the KCs of ID; *Clec4f-Cre* mice compared to their Healthy; *Clec4f-Cre* controls, and r-mS100A9 treatment lowers the expression of both, which, in the ID; Clec4f-Cre; r-mS100A9, are indistinguishable from those of the Healthy; *Clec4f-Cre* controls. Also, data shown in Fig. 1F indicate that the level of expression of CD206 and CD80 in the MoMΦ isolated from ID; *Clec4f-Cre*; r-mS100A9 group is similar to ID; *Clec4f-Cre* mice and their Healthy; *Clec4f-Cre* controls. Collectively, these results indicate that r-mS100A9 treatment rescues the polarization changes of KCs caused by ID.

To further characterize the beneficial action of S100A9 on KCs in ID, we administered ID; KC-INTACT mice with r-mS100A9 (ID; KC-INTACT; r-mS100A9 mice) (Fig. 2A). As expected, ID; KC-INTACT; r-mS100A9 mice displayed undetectable level of circulating insulin (Fig. S2D). Three hours after the r-mS100A9 injection, KC nuclei were isolated (as previously described^14,42^) and utilized for RNA-seq analysis. The data gathered from ID; KC-INTACT; r-mS100A9 mice were compared with the data obtained from ID; KC-INTACT (Fig. S2F) and Healthy; KC-INTACT (Fig. S2G) groups. Acute treatment with r-mS100A9 partially reverts the ID-induced expression abnormalities in 131 genes, with 93 of these genes showing RNA levels restored to a healthy state (Figs. 2B, E). Functional enrichment analysis revealed that 72% of the genes with altered expression in ID and rescued by r-mS100A9 administration are related to lipid metabolism (53% to triglyceride metabolic process, and 19% to fatty acid elongation), while other marginally enriched clusters were related to p53 signaling and cell cycle (Fig. 2F-G). mRNA levels of genes known to be involved in lipid uptake, trafficking, and storage (e.g., *Cyp2b10, Rab30, Acot4*)^50–52^ were significantly increased in KCs of ID; KC-INTACT mice as compared to Healthy; KC-INTACT controls (Fig. S2H-J). Also, mRNA levels of genes known to be involved in macrophage polarization switch and immune function (e.g., *Rorα*, *Cav1, Sgmgs2, Nifk*)^46,53–55^ were significantly altered in KCs of ID; KC-INTACT mice as compared to Healthy; KC-INTACT controls (Fig. S2K-N). Remarkably, mRNA levels of all the aforementioned genes were normalized by r-mS100A9 treatment as their contents were indistinguishable in KCs of ID; KC-INTACT; r-mS100A9 mice as compared to ID; KC-INTACT and Healthy; KC-INTACT groups (Fig. S2H-N).

To further analyze the transcriptional changes associated with macrophage polarization, we performed enrichment analysis for transcription factors known to regulate the transcription of genes whose expression is altered in ID; KC-INTACT and rescued in ID; KC-INTACT; r-mS100A9 mice. Among the most enriched transcription factors identified, several, such as PPARα, PPARγ, RORα, STAT3, GATA3, EGR1, BCL6, and others (Table S1), are reported to control polarization switch in macrophages^44,45,56–63^.

Collectively, our data show that r-mS100A9 treatment rescues polarization defects and lipid-related transcriptional alterations of the KCs caused by ID.

### S100A9 modulates the KC proteome, promoting an anti-inflammatory and lipid-homeostatic signature in ID

Our previous work shows that S100A9 normalizes diabetic hyperketonemia by acting *via* the TLR4 on hepatic non-parenchymal cells^38^. However, the specific cell type(s) in which TLR4 serves as the primary target of S100A9 is unknown. Based on the aforementioned data shown herein, we hypothesized that S100A9 acts on the KCs through TLR4 to normalize hyperketonemia induced by ID. Additionally, we directly tested whether the transcriptomic changes in KCs brought about by r-mS100A9 treatment in ID translated into proteomic changes. To this end, we performed label-free quantitative proteomic analysis of KCs isolated from mice expressing the TLR4 receptor only in KCs. We crossed mice carrying a loxP-flanked transcription blocking sequence between the second and third exon of the gene encoding for *Tlr4*, the *Tlr4^LoxTB^* allele^64^, with the previously mentioned *RIPDTR* mice, obtaining *Tlr4^+/+^; RIP-DTR* mice (*Tlr4^WT^*), *Tlr4^+/LoxTB^; RIP-DTR* mice (*Tlr4^HET^*), and *Tlr4^LoxTB/LoxTB^; RIP-DTR* mice (*Tlr4^KO^*) (Fig. S3A)^38^. The *Tlr4^LoxTB^* allele allows the Cre-conditional reactivation of the expression of the TLR4 receptor in specific cells and/or tissues. To accomplish KC-specific re-expression of TLR4 in ID mice, we crossed the *Tlr4^KO^* mice with the *Clec4f-Cre-tdTomato* mouse strain^41^, and we achieved re-expression of the TLR4 receptor only in the KCs (*Tlr4^KO^; RIP-DTR; Clec4f-Cre* mice, referred to as *Tlr4^KC^*) (Fig. 3A, S3A). Compared to their wild-type counterparts, *Tlr4^KO^* mice displayed undetectable levels of *Tlr4* mRNA in the liver, SkM, and adipose tissue (Fig. 3B). *Tlr4^KC^*mice displayed a partial, liver-specific rescue of *Tlr4* levels (Fig. 3B), in line with the expected re-expression of *Tlr4* in a fraction corresponding approximately to 15-20% of hepatic cells. We further validated the KC-specific re-expression of *Tlr4* by separating the KCs from the hepatocytes of *Tlr4^WT^*, *Tlr4^KO^*, and *Tlr4^KC^* mice (see methods for details). The identity of the two cell fractions was confirmed by the respective expression of *Albumin*, a marker of the hepatocytes, and the macrophage marker *F4/80* in the KCs (Fig. 3C). In the hepatocyte fraction, *Tlr4* mRNA content was similarly and significantly reduced in *Tlr4^KO^* and *Tlr4^KC^* compared to *Tlr4^WT^*mice (Fig. 3D). In the macrophage fraction, *Tlr4* mRNA content was significantly reduced in *Tlr4^KO^* and partially rescued in *Tlr4^KC^* compared to *Tlr4^WT^* mice (Fig. 3D). To further validate our model, we took advantage of the previously described *RIPDTR; Clec4f-Cre* mice (carrying the *Tlr4* wild-type allele). To test whether the Cre recombinase-driven recombination on the *Tlr4* allele was achieved only in the KCs, KCs from *RIPDTR*; *Clec4f-Cre* and *Tlr4^KC^* mice were sorted based on the TdTomato expression and these cells were separated from the other CD45+ immune cells (CD45 being a well-characterized marker of all the immune cells^12^) and CD45-cells (the remaining fraction containing contaminating parenchymal cells and non-immune non-parenchymal cells) (see methods section for details). The genotyping of the different cell fractions from *RIPDTR*; *Clec4f-Cre* and *Tlr4^KC^* mice confirmed the successful Cre recombinase-driven recombination only in the KCs; indeed, the Cre-recombinase-deleted *Tlr4^LoxTB^*allele was only present in KCs from *Tlr4^KC^* mice (Fig. S3A, B). Collectively, these results indicate that *Tlr4^KC^* mice express TLR4 only in KCs. *Tlr4^KC^* mice were rendered ID through DT treatment and then administered with saline (ID; *Tlr4^KC^*) or with the r-mS100A9 protein (ID; *Tlr4^KC^*; r-mS100A9) (Fig. 3A). ID; *Tlr4^KC^*and ID; *Tlr4^KC^*; r-mS100A9 mice displayed a similar degree of severe hypoinsulinemia (Fig. S3C). Three hours after saline or r-mS100A9 administration the KCs were isolated and assessed through Liquid Chromatography (LC) - Mass Spectrometry (MS) analysis (LC-MS/MS). Through label-free proteomic analysis, we quantified a total of 3359 proteins, of which 130 were significantly upregulated and 195 downregulated in ID; *Tlr4^KC^*; r-mS100A9 compared to their ID; *Tlr4^KC^* controls (Figs. 3E, F). We performed STRING analysis for GO annotations and KEGG pathways on differentially abundant proteins. r-mS100A9 promotes the expression of proteins involved in angiogenesis, macrophage polarization, immune cell recruitment, and lipid homeostasis^65–68^, while reducing or preventing the expression of proteins related to glucose and amino acid metabolism^69–72^, known to be pathologically enhanced in ID liver^33^.

**Fig. 3.**
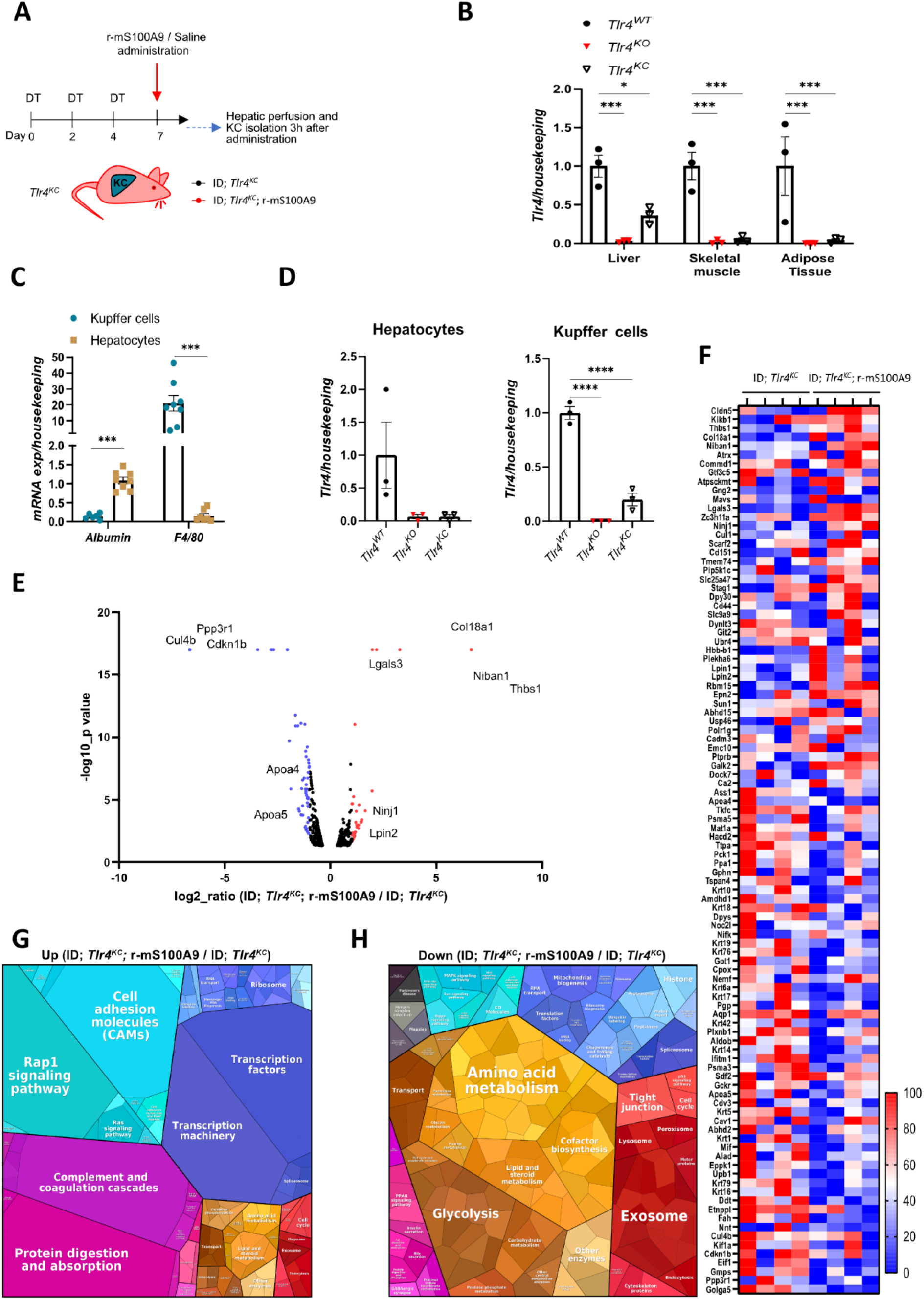
S100A9 modulates the KC proteome, promoting an anti-inflammatory and lipid-homeostatic signature in ID. **(A)** Schematic representation of experimental groups and treatment. Age-and gender-matched *Tlr4^KC^* mice received three DT injections on day 0, 2, and 4 to reach ID. On day 7, saline or r-mS100A9 were administered to these mice. 3h after, data and tissues were collected, and the KCs fraction was isolated from the liver for proteomic analysis (LC-MS/MS) and label-free protein quantification. **(B)** *Tlr4* mRNA content in the liver, skeletal muscle (gastrocnemius), and adipose tissue (interscapular brown adipose tissue, iBAT) of age-and gender-matched Tlr4^WT^ (n=3), Tlr4^KO^(n=3), Tlr4^KC^ (n=3) mice. **(C)** *Albumin* and *F4/80* mRNA content in the isolated KCs and hepatocytes of mice shown in (**B**). **(D)** *Tlr4* mRNA content in KCs and hepatocytes isolated from the indicated groups. **(E)** Volcano plot representing proteomic data from label-free protein quantification. Proteins are ranked according to their statistical p-value (-log10_p-value) and their relative abundance ratio (log2-ratio (ID; *Tlr4^KC^*; r-mS100A9/ ID; *Tlr4^KC^*)) between ID; *Tlr4^KC^;* r-mS100A9 and ID; *Tlr4^KC^* samples. In red, proteins with Fold Change≥2; in blue, proteins with Fold Change≤0.5. Names of selected metabolically relevant proteins are highlighted. **(F)** Heatmap showing the significant (p<0.05) mostly changed proteins (Fold change≥ or ≤2) detected in at least 3 samples. Each row is normalized from 0 to 100%. **(G-H)** Proteomaps illustrating functional changes in the KCs proteome upon r-mS100A9 treatment: treemap of proteins increased (fold change≥2) (**G**) or decreased (fold change≤0.5) (**H**) by r-mS100A9 treatment. Proteins with similar functions are classified according to KEGG orthology. Neighboring areas correspond to related functional groups and are marked by similar colors. The size of each polygon depicts the mass fraction of the respective proteins, reflecting protein abundances adjusted for their size. Error bars represent SEM, statistical analyses were done using one-way or two-way ANOVA (Tukey’s post-hoc test). *p ≤ 0.05, **p ≤ 0.01, ***p ≤ 0.001.

Among the proteins most increased by r-mS100A9 or detected only in the r-mS100A9-treated group (NIBAN1, THBS1, ATRX, ATPSCKMT, COMMD1, GTF3C5, KLKB1, CLDN5, COL18A1), several are involved in lipid metabolism, macrophage polarization, and inflammation (Fig. 3G, Fig. S3D)^67,68,73–76^. Interestingly, this analysis also brought up several proteins, whose expression was significantly increased or decreased by r-mS100A9, that have been associated with an anti-inflammatory signature of macrophages, such as NIBAN1, GTF3C5, GNG2, GALECTIN3, CDKN1B, and FBP1^77–82^ (Fig. 3F).

Among the proteins strongly downregulated by r-mS100A9 treatment or detected only in the saline-treated group (EIF1, CUL4B, PPP3R1, GMPS, GOLGA5, CDKN1B, KIF1A), several have been previously characterized in the context of metabolism and immunity^83–85^ (Fig. 3H, S3E). r-mS100A9 downregulates proteins involved in amino acid metabolism (GOT1, PAH, ASS1, UPB1, ALAS1, TDO2, FTCD, UROC1), some of which, such as Ass1,

(Fig. 3H, S3E) are already known to be upregulated in the liver in ID^33^. Moreover, r-mS100A9 downregulates proteins involved in gluconeogenesis (MDH1, TPI1, PGAM1, PGK1, ENO1, PCK1, FBP1, PGM1) (Fig.. 3F, S3E) and the amount of the other apolipoproteins involved in lipoproteins association and triglyceride transport (APOA1, APOC4, APOA4, and APOA5)^86^ is consistently reduced in the r-mS100A9 - treated group.

Overall, our proteomic analysis confirms that r-mS100A9 reprograms KCs in ID, promoting an anti-inflammatory and lipid-regulating phenotype while suppressing pathological metabolic processes. These findings underscore its therapeutic potential in ID.

### S100A9 rescues diabetic hyperketonemia and hypertriglyceridemia via TLR4 in KCs

In the context of ID, S100A9 exerts hyperketonemia-normalizing action *via* TLR4 expressed in hepatic non-parenchymal cells^38^. Thus, to directly test whether the beneficial effect of S100A9 on KC polarization/transcriptomic/proteomic aberrancies caused by ID is of pathophysiological relevance, we assessed whether TLR4 in the KCs is sufficient to mediate the S100A9-induced hyperketonemia-improving action in ID. To this end, *Tlr4^WT^*, *Tlr4^KO^*, and *Tlr4^KC^* mice were rendered ID through DT treatment and then underwent hydrodynamic tail vein injection (HTVI) of a plasmid driving the over-expression (OE) of *S100a9* under the control of the *Albumin* promoter (pLive_*S100a9*), as previously described^37,38^. We obtained the 5 groups represented in Fig. 4A: i) ID; *Tlr4^WT^*; S100A9^OE^, ii) ID; *Tlr4^KO^*; S100A9^OE^, and iii) ID; *Tlr4^KC^*; S100A9^OE^ and two additional groups of *Tlr4^WT^* mice as controls: iv) healthy; *Tlr4^WT^* (referred to as Healthy), and v) ID; *Tlr4^WT^*hydrodynamically injected with the control pLive empty plasmid (referred to as ID; *Tlr4^WT^*). Upon DT treatment, all the experimental groups reached a similar degree of ID and significant hyperglycemia compared to Healthy controls (Fig. S4A, B). Successful overexpression of S100A9 was observed in the liver (Fig. S4C, D), with a trend of increase in the circulation (Fig. S4E), in all groups that underwent pLIVE-*S100a9* HTVI. Moreover, all the ID groups presented a similar reduction in body weight compared to Healthy mice, as expected (Fig. S4F). Taken together, these results validate our animal model.

**Fig. 4.**
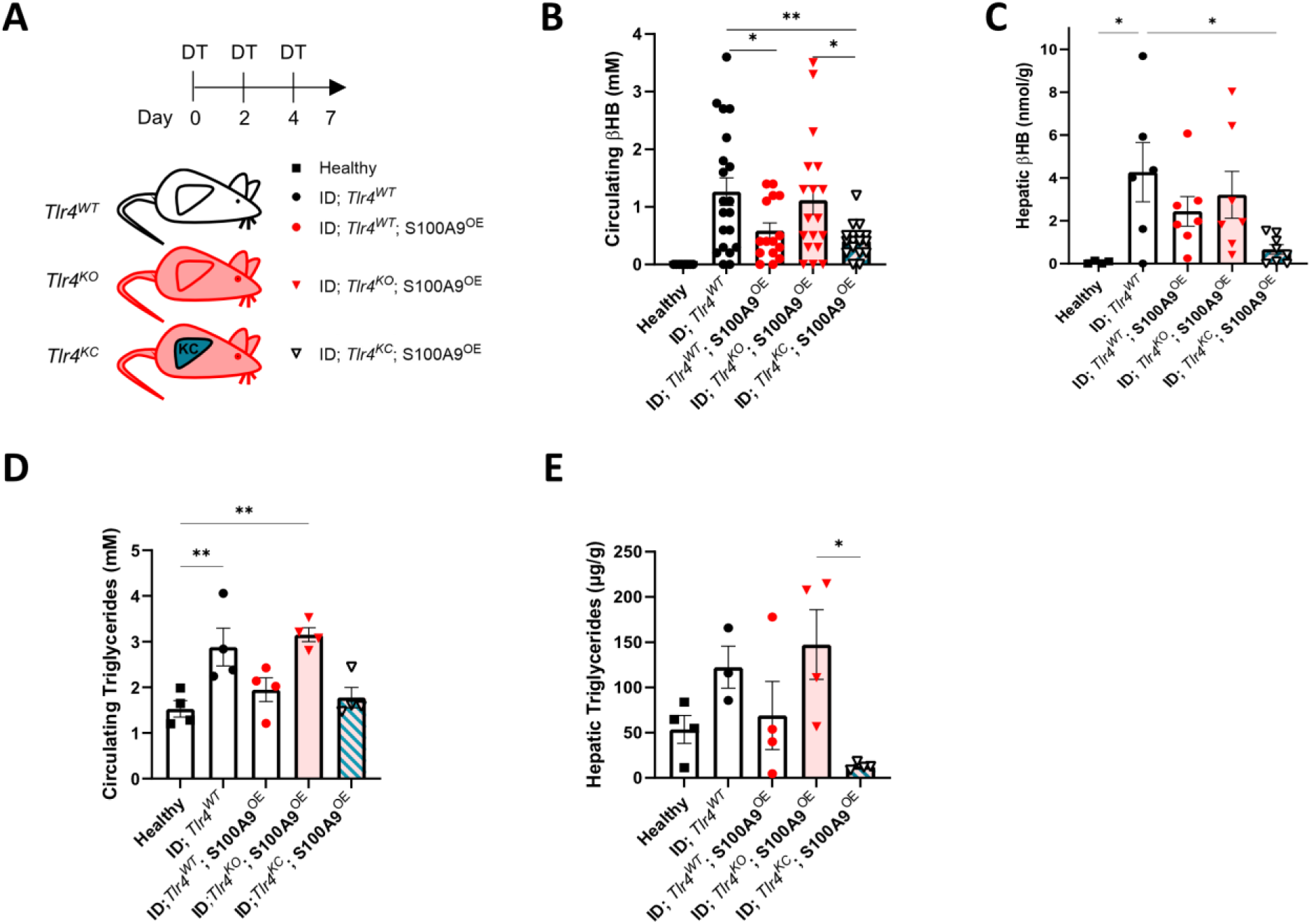
S100A9 rescues diabetic hyperketonemia and hypertriglyceridemia via TLR4 in KCs. **(A)** Scheme indicating the generation and treatment of the experimental groups: age-and gender-matched Healthy (n=4), ID; Tlr4^WT^ (n=19), ID;Tlr4^WT^;S100A9^OE^ (n=15), ID;Tlr4^KO^;S100A9^OE^ (n=18), ID;Tlr4^KC^;S100A9^OE^ (n=16). ID was achieved through 3 intraperitoneal DT injections at days 0, 2, 4, in *RIPDTR* mice. S100A9^OE^ was achieved through HTVI with pLive_*S100a9* plasmid on day 0, together with the first DT injection. Data were collected on day 7 after 3 h of fasting, and mice were euthanized. **(B-C)** Circulating βHB (**B**) and hepatic βHB levels (**C**) in the indicated cohorts on day 7. **(D-E)** Circulating (**D**) and hepatic (**E**) triglycerides in the indicated cohorts on day 7. Error bars represent SEM, statistical analyses were done using one-way or two-way ANOVA (Tukey’s post-hoc test). Comparisons were made with Healthy controls unless otherwise specified on the graph. *p ≤ 0.05, **p ≤ 0.01, ***p ≤ 0.001.

S100A9 overexpression counteracts the hyperketonemia caused by ID^37–39^. Indeed, circulating β-hydroxybutyrate (βHB) was elevated in ID;*Tlr4^WT^,* and similar in ID;*Tlr4^WT^*;S100A9^OE^ mice compared to Healthy mice (Fig. 4B). As previously shown, TLR4 is necessary for the S100A9-induced hyperketonemia-improving effect^38^. In keeping with these results, ID;*Tlr4^KO^*;S100A9^OE^ mice did not achieve the S100A9-induced hyperketonemia-improving action as they displayed a similar level of hyperketonemia as ID;*Tlr4^WT^*mice (Fig. 4B). Noteworthy, the hyperketonemia-lowering action of S100A9 was restored when TLR4 was re-expressed in the KCs. Indeed, the level of circulating βHB was similar between ID;*Tlr4^KC^*;S100A9^OE^, and ID;*Tlr4^WT^*;S100A9^OE^ mice (Fig. 4B). Data presented in Fig. 4C show that while the hepatic content of βHB is significantly increased in ID;*Tlr4^WT^* and ID;*Tlr4^KO^;*S100A9^OE^ mice, it tends to be elevated in ID;*Tlr4^WT^*;S100A9^OE^ and is similar (i.e., normalized) in ID;*Tlr4^KC^;*S100A9^OE^ mice as compared to healthy control (Fig. 4C).

S100A9 ameliorates hypertriglyceridemia caused by ID^37^. Yet, the underlying molecular and cellular mechanism is unknown. First, we assessed whether TLR4 is required for this effect. As expected, the circulating level of triglycerides was increased in ID;*Tlr4^WT^* mice compared to the Healthy controls (Fig. 4D). Mice overexpressing S100A9 in the context of intact TRL4 (i.e., ID;*Tlr4^WT^*;S100A9^OE^ mice) had improved hypertriglyceridemia as compared to ID;*Tlr4^WT^* mice; yet, this effect of S100A9 was blunted in the context of TRL4 deficiency (i.e., ID;*Tlr4^KO^*;S100A9^OE^ mice) (Fig. 4D). Next, we tested the relevance of TLR4 in KCs. As shown in Fig. 4D, mice expressing TLR4 only in KCs (ID;*Tlr4^KC^*;S100A9^OE^ mice) had similar circulating levels of triglycerides as ID;*Tlr4^WT^*;S100A9^OE^ mice and healthy controls. Similarly, hepatic triglyceride content was increased in ID;*Tlr4^WT^* and ID;*Tlr4^KO^*;S100A9^OE^ mice and normalized by S100A9 overexpression in ID;*Tlr4^WT^*;S100A9^OE^ and ID;*Tlr4^KC^*;S100A9^OE^ groups (Fig. 4E). These results reveal a previously unrecognized role of TLR4 in mediating the hypertriglyceridemia-lowering effect of S100A9 and demonstrate that TLR4 in the KCs underlies this action in ID.

In conclusion, S100A9’s ability to normalize hyperketonemia and hypertriglyceridemia in ID is primarily driven by TLR4 signaling in KCs, uncovering a novel mechanism for regulating metabolic dysfunction.

### S100A9 exerts glucoregulatory effects in SkM via TLR4 on immune cells

In addition to its effects on ketone and lipid metabolism, enhanced S100A9 also exerts glucose-lowering action in ID^39^. This glucoregulatory action is achieved *via* promoting SkM glucose uptake in a TLR4-dependent manner^39^. Yet, it is unknown in which cell type(s) TLR4 mediates this beneficial action of S100A9.

First, we began by testing the relevance of TLR4 in myocytes. To this end, an adeno-associated virus (AAV) expressing Cre-recombinase with a GFP tag was injected intra-muscularly in *Tlr4^KO^* mice (referred to in Fig. S5A as Cre-GFP SkM). As a control, intra-muscular injection of an AAV expressing GFP was performed in the contralateral leg of the same mice (referred to in Fig. S5A as GFP SkM), as mentioned above. Western blot assay revealed GFP content in SkM isolated from injected legs; yet, GFP protein content was absent in other tissues including the liver, interscapular brown adipose tissue (iBAT), perigonadal white adipose tissue (pWAT), and plasma of injected *Tlr4^KO^*mice (Figs. S5A-B). Polymerase chain reaction (PCR) and quantitative PCR (qPCR) analysis confirmed amplicons corresponding to the size of the Cre-recombinase-modified *Tlr4^LoxTB^* allele and regain of *Tlr4* transcript that was present only in the SkM from the Cre-recombinase injected leg and absent from the contralateral leg and aforementioned tissues (Fig. S5C-D). To determine if Cre-recombinase expression was restricted to myocytes, we separated them from non-myocyte cell fractions from digested SkM. Western blot for the myocytes marker Troponin I^87^ and GFP confirmed robust expression in injected whole muscle that was absent from the non-myocyte cell fractions (Fig. S5E). PCR also confirmed the absence of the Cre-recombinase-modified *Tlr4^LoxTB^* allele in the fractions of Cre-recombinase injected SkM (Fig. S5F).

We then assessed the effect of myocyte-specific regain of TLR4 on S100A9-mediated glucose uptake by generating the groups shown in Fig. 5A. Once again, r-mS100A9 treatment slightly improved glycemia and enhanced SkM glucose uptake in TLR4 intact mice (i.e., ID; Tlr4^WT^; r-mS100A9 mice), which was lost in ID; *Tlr4^KO^;* r-mS100A9 mice (Fig. 5B-C). Strikingly, regain of TLR4 in myocytes of the Cre-recombinase injected leg in ID;Tlr4^Myo^; r-mS100A9 mice failed to improve hyperglycemia and increase glucose uptake (Fig. 5B-C) indicating that myocyte TLR4 is dispensable for S100A9’s glucoregulatory effects.

**Fig. 5.**
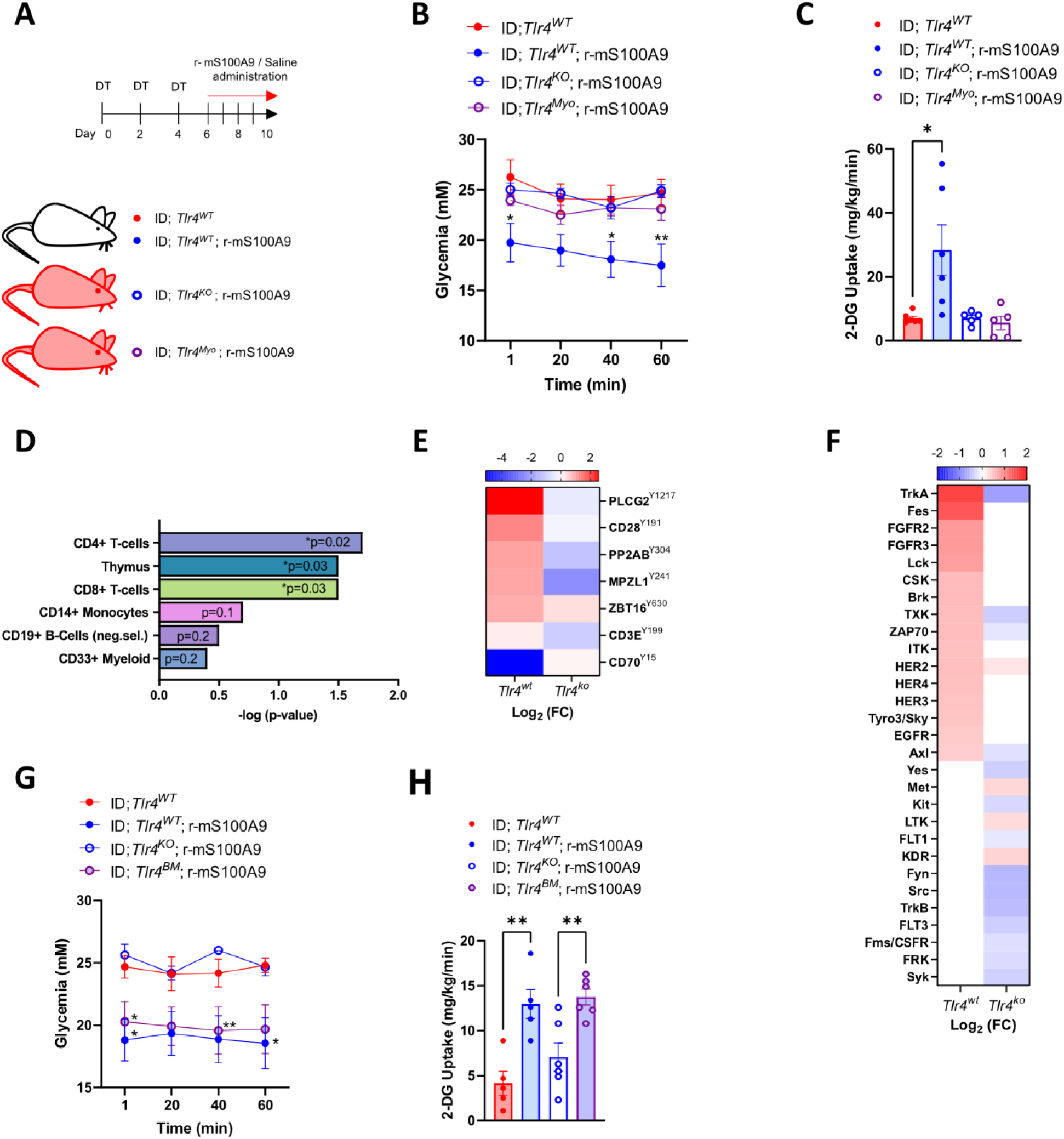
S100A9 exerts glucoregulatory effects in skeletal muscle *via* TLR4 on immune cells. **(A)** Scheme indicating the generation and treatment of indicated experimental groups (n= 5-6 per group). A week before the first DT injection, *Tlr4^Myo^*mice received an intra-muscular injection in the hind limb of 1×10^9 PFU AAV expressing Cre-GFP, while *Tlr4^KO^* mice received the same virus with only GFP. ID was achieved by three intraperitoneal DT administrations (at days 0, 2, and 4) in *RIP-DTR* mice. Starting from day 6, mice were administered twice daily with either 50 μl of saline or r-mS100A9. Metabolic parameters were followed over 4 consecutive days of treatment. **(B-C)** Glycemia (**B**) and 2DG glucose uptake (**C**) in isolated soleus muscle from indicated groups 1 hour after a bolus of 2DG. **(D-F)** EnrichR analysis of predicted cell type (**D**), Log2 of fold change (FC) of most differentially changed phospho-sites (**E**) and predicted kinases (**F**) in ID; *Tlr4^KO^* and ID; *Tlr4^WT^* soleus muscle in response to r-mS100A9 treatment. **(G-H)** Glycemia (**G**) and 2DG glucose (**H**) uptake 1 hour after a bolus of 2DG from ID; *Tlr4^WT^* (n=5), ID; *Tlr4^WT^*; r-mS100A9 (n=5), ID; *Tlr4^KO^;* r-mS100A9 (n=6), ID; *Tlr4^BM^;* r-mS100A9 (n=6) treated mice. Error bars represent SEM, and statistical analyses were done using one-way ANOVA or two-way ANOVA (Tukey’s post hoc test) when more than two groups and more than one experimental condition/time point were compared. * indicates comparison to ID; *Tlr4^WT^* controls unless otherwise indicated on the graph. *P < 0.05, **P < 0.01.

To gain further insights into how S100A9 is promoting glucose uptake, we performed a protein tyrosine kinase profiling (PamGene^88^) on whole muscle lysates of ID; *Tlr4^WT^* and ID; *Tlr4^KO^* mice treated with saline or r-mS100A9. Upstream kinase assay predicted several differentially regulated kinases in ID; *Tlr4^WT^* and ID; *Tlr4^KO^* SkM in response to r-mS100A9. Enrichment analysis^89^ results of these differentially regulated kinases based on cell type indicated the involvement of immune cells, notably CD4+ and CD8+ T-cells, as well as monocytes, B cells, and CD33+ myeloid cells (Fig. 5D). Ranking of the top most differentially phosphorylated phosphosites (Fig. 5E) and predicted kinases (Fig. 5F) additionally reveal many kinases and phosphosites involved in T-cell function (Lck, CSK, TXK, ZAP70 and ITK)^90^ and immune cell function, such as Phospholipase C-gamma 2 (PLCγ2)^91,92^ and tropomysin receptor kinase A (TRKA)^93,94^.

TLR4 is broadly expressed in myocytes as well as various non-parenchymal cells, including immune cells^95,96^. Considering the aforementioned data, we tested whether TLR4 in immune cells in SkM underlies the glucoregulatory action of S100A9.

To this end, we performed a bone marrow (BM) transplant whereby BM from *Tlr4^WT^* mice was isolated and injected into lethally irradiated *Tlr4^KO^* recipients to generate mice that have intact TLR4 in all hematopoietic stem cell (HSC)-derived immune cells^97^ (*Tlr4^BM^* mice). PCR from immune (CD45+) cells from digested SkM generated amplicons corresponding to the *Tlr4^WT^* allele that was absent from other non-immune (CD45-) cells and myocytes which displayed amplicons corresponding to the *Tlr4^LoxTB^* allele (Fig. S5G). We then subjected mice to the same treatment regimen as outlined in Fig. 5A and tested the effect of r-mS100A9 on SkM glucose uptake. Strikingly, regain of immune cell TLR4 was sufficient to restore S100A9’s glucose lowering effect and uptake into SkM (Fig. 5G-H). Taken together, our results demonstrate that immune cell TLR4 is sufficient for r-mS100A9 to lower hyperglycemia and enhance SkM glucose uptake. We then tested if these alterations in glucose metabolism have any effect on r-mS100A9’s dyslipidemia improving effects by measuring plasma ketones and triglycerides in the aforementioned groups. Circulating ketones and triglycerides were once again lowered by r-mS100A9 in a TLR4 dependent manner (Fig. S5H-I). However, this effect was not observed in mice expressing TLR4 only in HSC derived immune cells (Fig. S5H-I). These results indicate that the lipid and ketone lowering effect of S100A9 are dissociated from its glucose lowering effect and occur *via* TLR4 in distinct immune cell niches.

## Discussion

In this study, we unmasked immune niches within SkM and liver as putative targets for improving i) glucose imbalance and ii) ketone/lipid metabolic derangement caused by ID, respectively. Mechanistically, we unravelled that while TLR4 in BM-derived immune cells mediates the hyperglycemia-improving effect of r-mS100A9 within the SkM, TLR4 in KCs underlies the hyperketonemia-and hypertriglyceridemia-lowering action of S100A9 in the liver. This conclusion was drawn mainly because our results shown in Fig. 5B-C indicate that re-expression of TLR4 in SkM of ID mice is unable to mediate the hyperglycemia-improving and glucose-uptake-enhancing action of S100A9. However, data shown in Figs. 5G-H suggest that ID mice lacking TLR4 in all cells except for BM and BM-derived immune cells were able to properly respond to the aforementioned glucoregulatory actions of S100A9. While our data clearly suggest that TLR4 in myocytes is not required (Fig. 5B-C), the precise identity of immune cell type(s) in which TLR4 underlies S100A9-induced SkM glucose uptake remains to be elucidated. Yet, the predicted cell types and the differentially phosphorylated sites (notable CD28, a well-known T cell co-receptor^98^), emerged in the Pamgene analysis of the SkM from r-mS100A9-treated ID mice, pinpoint the T cells as top candidates to mediate the effect of r-mS100A9 in this tissue (Figs. 5D-F). Thus, further investigations aimed at manipulating TLR4 exclusively in T cells are warranted. The conclusion that TLR4 in the KCs mediates the ketone/lipid improving action of S100A9 is strongly supported by the fact that re-expression of TLR4 in KCs of ID mice is sufficient for virtually all of the hyperketonemia-and hypertriglyceridemia-lowering effect of S100A9 (Figs. 4B-E). Moreover, several studies have shown that in adulthood KCs do not originate from bone marrow-derived immune cells (unless following a major hepatic injury)^12,41,99^. Hence, our data shown in Fig. S5H-I indicating that ID mice expressing TLR4 only in BM and BM-derived immune cells do not respond to the hyperketonemia-and hypertriglyceridemia-lowering actions of S100A9, further corroborate the notion that TLR4 in KCs underpins the ketone and lipid improving effect of S100A9.

Untreated ID is a deadly catabolic condition^33^. Interestingly, ID shares several features of a starvation state, as, for example, enhanced ketogenesis, proteolysis, and lipolysis, accompanied by reduced body weight ^33,100^. While the changes in hepatic immune landscape induced by starvation are unknown, it is tempting to speculate that this change might share some similarities with the ones shown here in the context of ID (Fig. 1C). However, this appears unlikely. For example, reduced energy intake is associated with dampened hepatic inflammation ^14^. However, data shown in Figs. 1E, 2B-G indicate that in ID KCs shift towards what has been defined as a pro-inflammatory KC2 phenotype (identified as a CD206^HIGH^ KC subset)^9^. These results align with our previously published data showing increased tissue and systemic inflammation in ID mice^39^. The KC2 gene signature is also associated with lipid metabolism and lipid-associated oxidative stress, and a recent study shows an increased presence of the KC2 population in obese mice^9,12^. Hence, ID impacts KCs polarization in a way that is more similar to states of nutrient excess (e.g. obesity)^9^ which do not display enhanced ketogenesis ^15^. Nevertheless, S100A9 suppresses ID-induced ketogenesis and at the same time reverts the KCs polarization towards a healthier KC1 phenotype (Figs. 1E, 4B-C). These data are also in agreement with our previously published data showing the potent anti-inflammatory effect of S100A9 in ID mice^39^. Interestingly, the results presented here raise the question about the modality by which i) ID negatively affects KCs polarization/activity and ii) S100A9-induced change of KCs polarization/activity leads to suppression of enhanced ketogenesis, a metabolic process that occurs in hepatocytes^34^. While the mechanism by which ID alters KCs polarization/activity is unknown, our KC proteomic data shown in Fig. 3E-H suggest that several proteins affected by S100A9 in ID may contribute to the crosstalk between KCs and hepatocytes. For example, S100A9 increases the amount of thrombospondin 1 (THBS1), kallikrein (KLKB1), and galectin 3 (GALECTIN3) in KCs of ID mice (Figs. 3E-F, S3D). The role of these proteins in hepatic lipid metabolism is partially known^67,75,77^, whether they regulate ketogenic pathways is still unknown. Therefore, it is plausible that one, or more, of these proteins could be the S100A9-induced signal emanating from the KCs that instructs hepatocytes to suppress diabetic ketogenesis. To tackle this point, the metabolic characterization of genetically-engineered ID animal models overexpressing one, or more, of the aforementioned lead candidates specifically in KCs will be needed. These studies could also elucidate, at least in part, the mechanism(s) by which TLR4 in KCs underlies the hypertriglyceridemia-lowering action of S100A9.

Taken together, our results demonstrate that S100A9 is an efficient tool to target immune niches in different tissues to improve ID-induced metabolic derangements (hyperglycemia *via* the immune cells in the muscle, and hypertriglyceridemia and hyperketonemia *via* KCs in the liver) by TLR4-dependent mechanism(s).

## Methods

### Animal studies

All animal studies adhered to the ethical regulations of the University of Geneva, within the procedures approved by the animal care and experimentation authorities of the Canton of Geneva, Switzerland (animal protocol number GE/112A). Mice were kept with a standard chow diet and water available ad libitum in a temperature-controlled environment with a 12h light/dark cycle. All the experimental mice used for the study were 8-12 weeks old, with a weight between 25 and 30g, and had a mixed genetic background. To induce ID we utilized *RIPDTR* mice as previously characterized^37^. Healthy mice were intraperitoneally injected with Diphtheria Toxin (DT, Sigma Aldrich) dissolved in 0.9% NaCl (1.5μg/kg) three times on days 0, 2, and 4. Metabolic assessments were conducted on day 7 after the first DT injection. The *Tlr4* null allele (*Tlr4^LoxTB^*) was generated as previously documented ^64^. Briefly, the allele has a LoxP-flanked transcription blocking (TB) cassette inserted in the sequence between the second and the third exon of the gene encoding for the TLR4 receptor. Mice carrying the *Tlr4^LoxTB^*allele were bred with *RIPDTR* mice, obtaining *Tlr4^+/+^;RIPDTR* mice (*Tlr4^WT^*), *Tlr4^LoxTB/+^;RIPDTR* mice (*Tlr4^HET^*), and *Tlr4^LoxTB/LoxTB^;RIPDTR* mice (*Tlr4^KO^*)^38^. The *Tlr4^LoxTB^*allele allows the Cre-conditional reactivation of the expression of the TLR4 receptor in specific cells and/or tissues (*Tlr4 wild type* Forward F’ CTGACTGGTGTGAAGTGGAATATC, *Tlr4 LoxTB* Forward F2’ GTCATAGATGCATGCCAGATACA, *Tlr4* Reverse R’ CTGGACAAACAGTGGCTGGA primers were used for genotyping these animals, Fig. S3A). By crossing the *Tlr4^KO^*mice with the *Clec4f-Cre-tdTomato* mouse strain (Jax strain 033296, previously described^41^), we achieved the re-expression of the TLR4 receptor only in the KCs (*Tlr4^KC^*). The overexpression of S100A9 was achieved according to previous studies ^37,38^. Concisely, experimental mice were injected through HTVI with 50μg of a plasmid pLIVE-*S100a9* or with a pLIVE empty control (Mirus) on the day of the first DT injection. pLIVE-*S100a9* drives the expression of *S100a9* under the control of the *Albumin* promoter, therefore in the liver. Mice receiving acute treatment with the r-mS100A9 were injected via tail vein with 0.6mg/kg of protein or an equal volume of saline as control, as previously done^38,39^. Simultaneously with the injection, the mice were put in fasting, and three hours after, metabolic assessments were performed and blood and tissues were collected for analysis. For chronic r-mS100A9 treatment, mice received a subcutaneous injection with 1.2mg/kg of r-mS100A9 or an equal volume of saline twice daily for 4 consecutive days, as previously done^38,39^. On day 10, mice were euthanized after 3h of fasting, and the liver was collected for FACS analysis. *CAG-Sun1/sfGFP* mice (Jax strain 021039, already characterized ^101^) used in this study were crossed with *Clec4f-Cre-tdTomato; RIPDTR* mice to generate *KC-INTACT* mice with KC-specific GFP tagging of the nuclear membrane, plus the *RIPDTR* gene. *KC-INTACT* mice were rendered ID through 3 DT injections on days 0, 2, and 4, and on day 7 ID mice were acutely injected with saline or r-mS100A9 (0.6mg/kg). After 3h the experimental mice were euthanized and GFP-tagged nuclei were purified for RNA-seq analysis.

### Assessment of glucose uptake

In vivo glucose uptake in tissues was determined by a 10 μCi bolus retro-orbital injection of [^14^C]2-deoxyglucose which was administered 2 hours after subcutaneous injection (0.6 mg/kg) of r-mS100A9 or saline. Blood sampling was regularly done for the determination of 2DG kinetic in systemic blood, and after 1h mice were rapidly euthanized by cervical dislocation, and tissues were removed and stored at −80°C until use.

### Bone Marrow Transplants

Bone Marrow Transplants were performed as previously described^102^. Briefly, 8-week-old recipient *Tlr4^KO^* mice were lethally irradiated with 1,000 rad. 24 hours later, animals were reconstituted via retro-orbital injection of 5×10^6^ bone marrow hematopoietic cells collected from donor *Tlr4^WT^* mice. Mice were treated with Baytril 10% (6 mL/L drinking water) for the following 2 weeks. Five weeks after retro-orbital injection, the reconstitution was confirmed by analysing the presence of *Tlr4^WT^* allele in peripheral blood and immune cells isolated from SkM. Animals were subsequently kept ad libitum on a standard chow diet and subject to induction of ID and r-mS100A9 treatment as per the described protocol.

### Muscle cell isolation

Non-parenchymal and immune cell fractions of SkM were isolated as previously described^21,103^. Mice were euthanised and hind limb soleus muscles were immediately dissected and muscles were minced in a petri dish on ice using a surgical blade and 90 degree angled surgical scissors. Next, the minced muscle tissue was enzymatically digested in digestion buffer containing 2 mg/mL Dispase II (D4693, Sigma-Aldrich), 2 mg/mL Collagenase IV (17104019, ThermoFisher Scientific), 1 U/mL DNase (D4263, Sigma) and 2 mM CaCl2 in PBS at 37°C for 25 min, with gentle shaking every 5 min. The reaction was stopped by adding an equal volume of cold 20% FBS in HBSS and the suspension was passed through a series of 70μm cell strainers to remove tissue debris. Single cell suspensions were either subject to FACS analysis (as described), western blotting, or DNA extraction and PCR. Primary single suspensions of popliteal lymph nodes were prepared by gently straining the tissue through a 70μm cell strainer into FACS buffer (2% w/v BSA, 1mM EDTA in PBS).

### Intramuscular injection

Mice were anesthetized with isoflurane, and hind limbs were injected with 1x 10^9^ PFU of Adeno-associated virus serotype 9 (AAV9) expressing Cre recombinase and GFP, or GFP only, under the control of a Cytomegalovirus (CMV) promoter. The virus was dissolved in 30μl of saline. Seven days after injection, muscles were dissected for validation of Cre expression and recombination of the *Tlr4^LoxTB^* allele.

### Kinase activity profiling

Protein Kinase (PK) activity profiling was performed using “Pamgene Station”. A total of 5 µg of protein was extracted from soleus SkM using M-PER (Mammalian Protein Extraction Reagent, Thermo Fisher) and processed with PamChip4 microarray chips, provided by PamGene (PamGene International, the Netherlands) and printed at their facilities.

### Blood chemistry

Circulating glucose and βHB levels were assessed after 3h of fasting using a glucose and ketone meter (Statstrip Xpress, Nova Biomedical) on tail vein blood. Hormones and metabolite levels were measured in plasma. Plasma was obtained by centrifugation of tail vein blood in EDTA-coated tubes (Kent Scientific) (2000g x 10’), and essayed using commercially available kits: S100A9 (R&D systems, DY008), Insulin (CrystalChem, 90080), βHB (Sigma-Aldrich, MAK041), Triglycerides (DiaSys, 1 5700 99 10 030).

### Assessment of hepatic βHB, Triglycerides, and S100A9 content

10mg of liver were homogenized in lysis buffer (Tris 20 mM, EDTA 5 mM, NP40 1% (v/v), protease inhibitors (Sigma, P2714-1BTL)) and centrifuged at 17950g for 15 min at 4°C. For S100A9 measurements (R&D systems, DY008), the supernatant was then further diluted 1:50 in lysis buffer before ELISA analysis. Protein concentration in liver lysates was measured through Pierce BCA Protein Assay kit (ThermoFisher, 23225). For measuring hepatic triglycerides content, 100mg of liver was homogenized in 1.7mL of chloroform:methanol (1:1), centrifuged (2000g for 10 min at 4°C), and resuspended in 0.4mL of isopropanol, before quantification according to the kit’s manufacturer’s instruction (DiaSys, 1 5700 99 10 030).

### Kupffer cell isolation

KCs were isolated from the liver as previously described ^40,104^. In brief, mice were anesthetized using Ketamine/Xylazine, and a catheter was inserted into the portal vein. The liver was perfused with around 100mL washing solution (HBSS1X, EGTA 0.5mM) for 10min at speed 8 after the opening of the abdominal vena cava. After washing, the liver was perfused with 50mL digestion solution (HBSS1X, CaCl_2_ 1mM, Collagenase type 4 (Bioconcept, LS004188) 0.64mg/mL) for 5min at speed 8. After perfusion, hepatic cells were released by gently scraping the liver in 5mL HBSS1X-CaCl_2_ 1mM, the resulting solution was filtered through a 100μm cell strainer (Corning), further diluted in 15mL HBSS1X-CaCl_2_ 1mM, and centrifuged (50g for 3min at 4°C). The supernatant is layered on a Percoll gradient and centrifuged for 30min at 4°C 1200g without brakes. The middle interphase containing the KCs is transferred in a clean tube and washed twice with PBS, centrifuging for 10min 1500rpm at 4°C. The isolated KCs are then plated in complete Dulbecco’s Modified Eagle’s Medium (DMEM), 10% Fetal Bovine Serum (FBS), 1% penicillin and streptavidin. 30 min later, KCs were resuspended in TRIzol (Invitrogen) for RNA extraction or lysis buffer for protein analysis.

### GFP-tagged nuclei purification

Nuclei extraction was performed as previously described ^42^. In brief, 500mg of grinded liver tissue from KC-INTACT mice (GFP^KC^) was homogenized with a Dounce and a low pestle in 5mL low sucrose buffer (0.25 M sucrose, 25 mM KCl, 5 mM MgCl2, 20 mM Tris-HCl (pH 7.5), 1 mM 1,4-dithiothreitol (DTT), 0.15 mM spermine, 0.5 mM spermidine, 13 EDTA-free protease inhibitor cocktail (PIC) (Roche), and 60 U/mL RNasin Plus Rnase Inhibitor (Promega)), then added 0.35% Igepal CA-630 (Sigma), and further homogenized with the Dounce 5 times with a tight pestle. The homogenate was filtered through a 100μm filter (Corning), centrifuged 600g for 10 min, resuspended in 9X volume of high sucrose buffer (same as low sucrose buffer, but with 2M sucrose), and centrifuged again 15,000g for 15 min. The nuclei pellet was resuspended in wash buffer (as low sucrose buffer with 0.35% Igepal CA-630) and nuclei were pre-cleared through a 15min incubation with Protein G Dynabeads (Life Technologies). Beads were removed with the DynaMag2 magnetic rack (ThermoFIsher) and the nuclei were incubated with 3μg of anti-GFP antibody (ThermoFisher, G10362) for 30min. Thereafter, 80μL of Dynabeads were added to the sample, and incubated for 20min. Bead-bound nuclei were washed with wash buffer using the magnet. All steps were performed on ice or in a cold room at 4°C, and all the incubation steps on an end-to-end rotor. After the isolation of bead-bound nuclei, RNA was extracted for RNA sequencing analysis.

### Nuclei RNA isolation and sequencing

Bead-bound nuclei isolated from KC-INTACT mice were resuspended in TRIzol. After further purification steps in chloroform and ethanol, samples were loaded onto RNeasy MicroColumn (QIAGEN, 74004). RNA quantification was performed with a Qubit fluorimeter (ThermoFisher Scientific) and RNA integrity was assessed with a Bioanalyzer (Agilent Technologies). The Illumina Stranded Total RNA Prep, Ligation with Ribo-Zero Plus kit was used for the library preparation with 1 to 5 ng of total RNA (depending on availability) as input. Library molarity and quality were assessed with the Qubit and Tapestation (DNA High sensitivity chip, Agilent Technologies). Libraries were sequenced on a NovaSeq 6000 Illumina sequencer for SR100 reads. The FastQ files were mapped to the Ensembl GRCm39 mouse reference with STAR v2.7.10b ^105^. The differential expression analysis was performed with the statistical analysis R/Bioconductor package edgeR 1.38.4 ^106^. After normalization, the poorly detected genes were filtered out and out of 56’941 total genes, 17’175 genes with counts above 10 were kept for the analysis. To visualize the network and perform functional enrichment analysis and transcription factor enrichment, Heatmapper^107^ and Cytoscape ^108^ were used, together with the ClueGO ^109^, CluePedia, and iRegulon apps.

### FACS analysis

For FACS liver immunophenotyping mice were anesthetized and perfused as before (refer to Methods, Kupffer cell isolation). After perfusion, hepatic cells were released by gently scraping the liver in 5mL HBSS1X-CaCl_2_ 1mM, the resulting solution was filtered through a 100μm cell strainer, further diluted in 10mL HBSS1X-CaCl_2_ 1mM. 5 million cells were incubated in 4mL ACK lysis buffer (ThermoFisher, A1049201) for 10min in an end-to-end rotator, spun and resuspended with FACS buffer (PBS1X, BSA1%, EDTA 1mM). The sample was then incubated with viability dye (ThermoFisher, L34975, 1:200) for 20min according to the manufacturer’s instruction, and blocked with CD16/CD32 (eBioscience, 14-0161-82). Cells were then incubated with the primary antibodies diluted 1:200 for 30 min (F4/80 BD567201, TIM-4 Biolegend 130008, Ly6C BD560596, CD25 BD566120, CD45 Biolegend103155, CD11c BD563735, CD11b BD563553, CD4 BD564667, Ly6G BD612921, CD19 BD612971, MHCII BD 748708, CD8 BD612898, CD3 Invitrogen 25-0031-82, CD206 BD568806, CD80 BD560016, CD11b BD612800, F4/80 Biolegen422926, CD45 VP2311293), and treated with BD Transcription factor buffer set (Biosciences, 562574) according to the manufacturer’s protocol. Lastly, incubation with the primary antibody FoxP3 was performed (FoxP3 Invitrogen 35-5773-82) and cells were washed (PBS1X) and resuspended in FACS buffer.

### Assessment of mRNA and protein content

Mice were euthanized and tissues were quickly frozen in liquid nitrogen, and stored at-80°C. RNA was extracted using TRIzol (Invitrogen), and complementary DNA was synthesized using SuperScript II (Invitrogen) and analyzed by quantitative Real-Time PCR (RT-qPCR) with Power SYBR Green PCR Master Mix (Applied Biosystem, 4367569). mRNA contents were normalized to housekeeping genes (18S, β-actin and GAPDH) mRNA levels. Assays were performed using an Applied Biosystems QuantStudio 5 Real-Time PCR System machine. For qPCR the following primers were used: *Albumin* F’ TGCTTTTTCCAGGGGTGT and R’ TTACTTCCTGCACTAATTTGGCA, *sfGFP* F’ AGTGCTTCAGCCGCTACC and R’ GAAGATGGTGCGCTCCTG, *18s* F’ CGGAAAATAGCCTTCGCCATCAC and R’ ATCACTCGCTCCACCTCATCCT, *F4/80* F’ CTTTGGCTATGGGCTTCCAGTC and R’ GCAAGGAGGACAGAGTTTATCGTG, *Tlr4* F’ GTGTAGCCATTGCTGCCAAC and R’ TGAAGATGATGCCAGAGCGG, *Gapdh* F’ CAAGGTCATCCATGACAACTTTG and R’ GGCCATCCACAGTCTTCTGG, β*-actin* F’ CATCGTGGGCCGCTCTA and R’ CACCCACATAGGAGTCCTTCT. mRNA measurements have been repeated three times for each RT-qPCR analysis. Proteins were extracted by homogenizing the sample in lysis buffer (Tris 20 mM, EDTA 5 mM, NP40 1% (v/v), protease inhibitors (Sigma, P2714-1BTL)), then resolved by SDS-polyacrylamide gel electrophoresis, and transferred to a nitrocellulose membrane by electroblotting. Incubation with primary antibodies (Tubulin (Proteintech, 11224-1-AP) 1:5000, S100A9 (BosterBio, PB9678) 1:1000, GFP (Abcam, Ab6556) 1:1000, Troponin I (ThermoFisher, PA5-37897) 1:1000) was performed overnight at 4°C. The membrane was lastly incubated with a 1:10,000 secondary antibody solution (IRDye 800CW, 926-32211, IRDye 680RD, 926-68070) for 1h at room temperature, before detection with LICOR Odyssey Imaging System.

### Proteomics

For the proteomics analysis, KCs were isolated from *Tlr4^KC^* mice as described before, sample preparation and label-free protein quantification were performed as previously described with adjustments^110^. Protein samples were prepared using the phase-transfer surfactant method with minor modifications. Samples were reduced with 5 mM TCEP, alkylated with 20 mM iodoacetamide, and digested with Lys-C and Trypsin at a 1:50 ratio. Digested peptides were dissolved in 0.1% (v/v) TFA in 3% acetonitrile (v/v) and desalted using MonoSpin C18 columns (GL Sciences Inc., Japan). Subsequently, an equal volume of ethyl acetate was added, and samples were acidified with 0.5% trifluoroacetic acid (TFA). After centrifugation, the upper organic phase containing sodium deoxycholate was removed, and the remaining sample was kept at-80°C for 15 min. The lower aqueous phase containing digested tryptic peptides was dried using a centrifugal vacuum concentrator. Dry peptides were dissolved in 0.1% (v/v) formic acid in 2% (v/v) acetonitrile and peptide concentration was measured on a Nanodrop spectrophotometer (Thermo Scientific) before mass spectrometry measurement. Data were acquired in a data-dependent acquisition mode and analyzed using Proteome Discoverer v2.4. Identification was performed using the Uniprot mouse database with default settings. FDR was 0.01 at both the peptide precursor level and protein level. Cross-run normalization was done with the total peptide amount. Protein ratio calculation was performed using the pairwise ratio algorithm. Hypothesis testing was done using background-based t-test. GO annotations, enrichment analysis, and KEGG pathways information were assembled using Perseus ^111^, Proteomaps ^112^, and EnrichR ^89,113,114^.

### Immunohistochemistry analyses

Paraffin-embedded liver tissue 6μm sections were mounted on glass slides, paraffin was removed through serial passages of 5min in xylene and PBS with decreasing concentrations of ethanol (100%, 70%, 50%, and 0%). Liver tissue frozen in milestone Cryoembedding Compound (Milestone Medical, 51420) was cryosectioned at 10μm. Antigen-retrieval was performed by boiling for 30min at 95°C the slide in Na-citrate buffer (pH 6). Sections were washed three times in PBS for 10min, and then saturated in blocking solution (PBS1X, BSA1%, Goat serum 5%, TritonX100 0.2%) for 1h at room temperature. Subsequently, sections were incubated at 4°C overnight with the primary antibody solution (PBS1X, BSA1%, Goat serum 5%, and the primary antibody: GFP (ThermoFisher, G10362) 1:50, F4/80 (Cell Signaling, 71299) 1:100). Sections were then incubated for 1h at room temperature with the secondary antibody solution (DAKO Antibody diluent (Agilent, S0809), and the secondary antibody: Alexa 488 (Invitrogen, A27034) 1:400, Alexa 594 (Invitrogen, A21209) 1:400). Lastly, sections were mounted with Vectashield with DAPI (Vector laboratories, H1500-10), and visualized using Leica Stellaris5 confocal microscope.

### Production of recombinant murine S100A9

Recombinant murine S100A9 was synthesized as previously described ^39^. The cDNAs encoding murine S100A9 proteins were cloned into a pET29+ vector including a streptavidin tag and a maltose-binding protein tag followed by a tobacco etch virus (TEV) protease-cleavage site for removal of the tags. The vector was expressed in Escherichia coli BL21 DE3 (Invitrogen) cells. Protein expression was induced at an A600 of 0.6 by the addition of 1 mM isopropyl-β-d-thiogalactopyranoside. Cells were harvested after 16-hour incubation at 20°C by centrifugation at 5000g for 20 min. Bacterial cells were resuspended in 200 mM NaCl, 50 mM tris-HCl (pH 8.0), 3% glycerol, and 3 mM ß-mercaptoethanol (ME). Cell lysis was done by shear forces using a Microfluidizer LM20 set at 136,000 kPa at 4°C. Lysate was centrifuged for 35 min at 35,000g at 4°C. The supernatant containing the soluble protein was filtered using a 0.22-μm strainer. It was applied to a 5-ml His-Trap FF and washed. Protein was eluted with 15 ml of elution buffer [50 mM tris (pH 8), 200 mM NaCl, 450 mM imidazole, 3% glycerol, and 3 mM 2-ME]. Protein was slowly concentrated using 30-kDa MWCO AMICON concentrator and then treated with Strep-Tev protease and dialyzed overnight against 1.8 liters of dialysis buffer [20 mM tris-HCl (pH 8), 150 mM NaCl, 10 mM 2-ME, 0.5 mM EDTA, and 5% glycerol]. The next day, the protein was applied to a 5-ml Strep-Tactin XT. Next, unbound protein was collected, washed with 15 ml of dialysis buffer, collected, and pooled with unbound. Protein was slowly concentrated using a 10kDa MWCO AMICON concentrator. Concentrated protein was applied to a Superdex 200, 10/300 column at room temperature, equilibrated with isotonic phosphate-buffered saline buffer [137 mM NaCl, 2.7 mM KCl, 8 mM Na2HPO4, and 2 mM KH2PO4 (pH 7.4)].

## Statistical analysis

Data sets were analyzed for statistical significance using PRISM (GraphPad). A two-tailed unpaired Student’s t-test was applied for comparisons between two groups, while one-or two-way ANOVA with Tukey’s post-hoc test was used for comparisons involving more than two groups.

## Acknowledgments

We thank Dr Oscar Vadas, Léna Falconnet, and Rémi Visentin for the recombinant protein production at the Proteins and Peptides Facility of University of Geneva (https://www.unige.ch/medecine/ppr2p-platforms/), Cécile Gameiro, Lan Tran, and Grégory Schneiter for cytofluorimetric analysis at the Flow Cytometry core facility, and Mylène Docquier and Natasha Civic for the RNA-seq at the iGe3 Genomics Platform of the University of Geneva.

## Author contribution

Conceptualization: G.L., G.U., G.R., and R.C. Methodology: G.L., G.U., G.R., P.D.S.T., S.Z., F.F., M.C., F.V., C.V.-D., P.C., Y.W., A.W. Funding acquisition:

G.R., R.C. Writing - original draft: G.L., G.U. Writing-review, editing: G.L., G.U., G.R., R.C., P.D.S.T., S.Z., C.W., M.K., Y.W., P.C., C.V.-D.

## Materials & Correspondence

Any additional information required is available from the lead contact upon request.

## Data availability

RNA-seq data have been deposited at GEO at accession number: GSE292486 and proteomic data have been deposited at the ProteomeXchange at accession number: PXD062104.

## Supplementary Information

Figs. S1 to S5

Table S1

