## Supplemental Figures for "Harnessing distinct tissue-resident immune niches *via* S100A9/TLR4 improves ketone, lipid, and glucose metabolism"

**The PDF file includes:**

Figs. S1 to S5

Fig. S1.

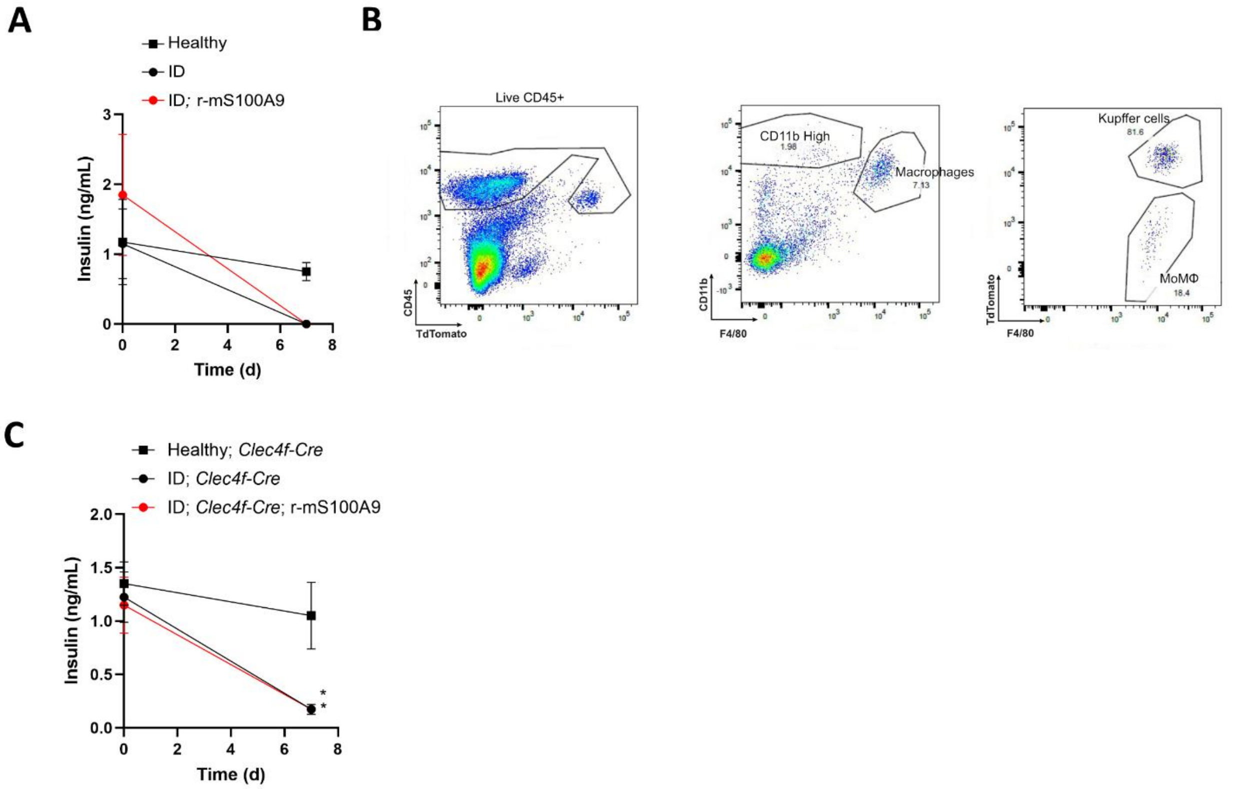


**Fig. S1. Insulin deficiency reshapes the immunophenotype of the liver.**

**(A)** Plasma insulin was measured on day 0 and 7 in Healthy, ID, and ID; r-mS100A9 mice as indicated in Fig. 1A.

**(B)** Flow cytometry plots showing the gating strategy used to identify different hepatic macrophage populations using the markers CD45, CD11b, TdTomato and F4/80.

**(C)** Plasma insulin was measured on day 0 and 7 in Healthy; *Clec4f-Cre*, ID; *Clec4f-Cre* and ID; *Clec4f-Cre*; r-mS100A9 mice as indicated in Fig. 1D.

Error bars represent SEM, statistical analyses were done using one-way or two-way ANOVA (Tukey's post- hoc test). Comparisons were made with Healthy controls. *$p \leq 0.05$.

**Fig. S2.**

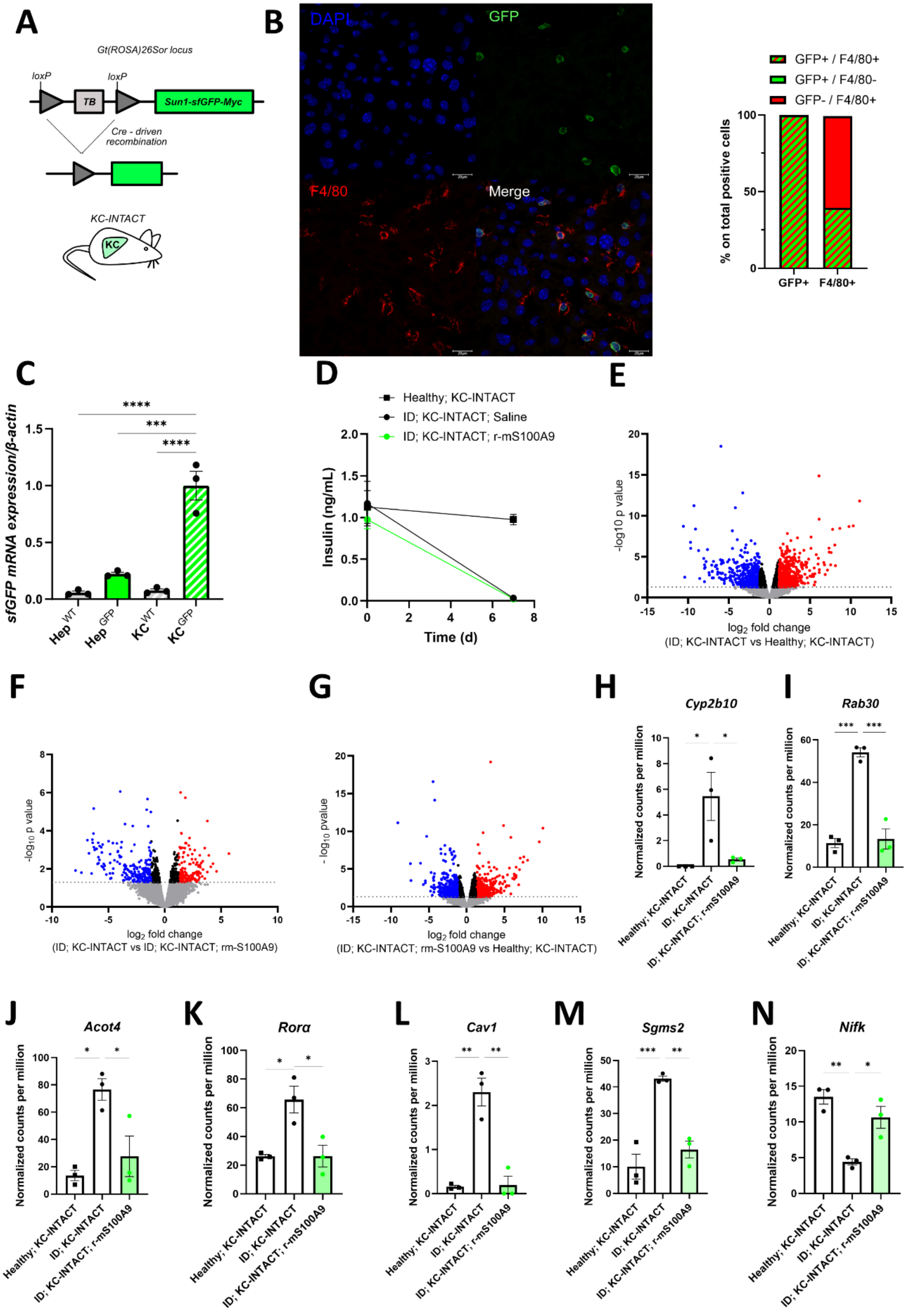
A
Gt(ROSA)26Sor locus
loxP
loxP
TB
Sun1-sfGFP-Myc
Cre - driven recombination
KC-INTACT
KC
B
DAPI
GFP
F4/80
Merge
GFP+ / F4/80+
GFP+ / F4/80-
GFP- / F4/80+
% on total positive cells
GFP+
F4/80+
C
sfGFP mRNA expression/β-actin
Hep WT
Hep GFP
KC WT
KC GFP
D
Healthy; KC-INTACT
ID; KC-INTACT; Saline
ID; KC-INTACT; r-mS100A9
Insulin (ng/mL)
Time (d)
E
-log10 p value
log2 fold change
(ID; KC-INTACT vs Healthy; KC-INTACT)
F
-log10 p value
log2 fold change
(ID; KC-INTACT vs ID; KC-INTACT; rm-S100A9)
G
- log10 pvalue
log2 fold change
(ID; KC-INTACT; rm-S100A9 vs Healthy; KC-INTACT)
H
Cyp2b10
I
Rab30
J
Acot4
K
Rorα
L
Cav1
M
Sgms2
N
Nifk
Normalized counts per million
Healthy; KC-INTACT
ID; KC-INTACT
ID; KC-INTACT; r-mS100A9

**Fig. S2. S100A9 rescues KC polarization and lipid-related transcriptional alterations caused by ID.**

**(A)** Scheme of the *Sun1-sfGFP* allele before (top) and after (bottom) Cre-recombinase-mediated deletion of the loxP-flanked transcriptional blocking (TB) sequences in KC-INTACT mice.

**(B)** Representative staining of GFP+ (green), F4/80+ (red), and DAPI+ nuclei in the liver of KC-INTACT mice. Scale bar=20μm (**left**). Average percentages of F4/80+ or GFP+ cells in the GFP+ or F4/80+, respectively, cell count calculated on 4 images from independent staining (**right**).

**(C)** *Gfp* mRNA content in the isolated KCs and hepatocytes from the control wild type ($Hep^{WT}$ and $KC^{WT}$) and KC-INTACT mice ($Hep^{GFP}$ and $KC^{GFP}$).

**(D)** Plasma insulin was measured on day 0 and 7 of the experimental groups shown in Fig. 2A.

**(E-G)** Volcano plot showing the $log_2$ fold change of ID; KC-INTACT versus Healthy; KC-INTACT (**E**), ID; KC-INTACT versus ID; KC-INTACT; r-mS100A9 (**F**), ID; KC-INTACT; r-mS100A9 versus Healthy; KC-INTACT (**G**) groups shown in Fig. 2A.

**(H-N)** Normalized counts per million of selected genes from the RNA-seq analysis of samples shown in Fig. 2B: *Cyp2b10* (**H**), *Rab30* (**I**), *Acot4* (**J**), *Rorα* (**K**), *Cav1* (**L**), *Sgms2* (**M**), *Nifk* (**N**).

Error bars represent SEM, statistical analyses were done using one-way or two-way ANOVA (Tukey's post- hoc test). Comparisons were made with Healthy controls, unless indicated otherwise on the graph. $*p \leq 0.05$, $**p \leq 0.01$, $***p \leq 0.001$.

**Fig. S3.**

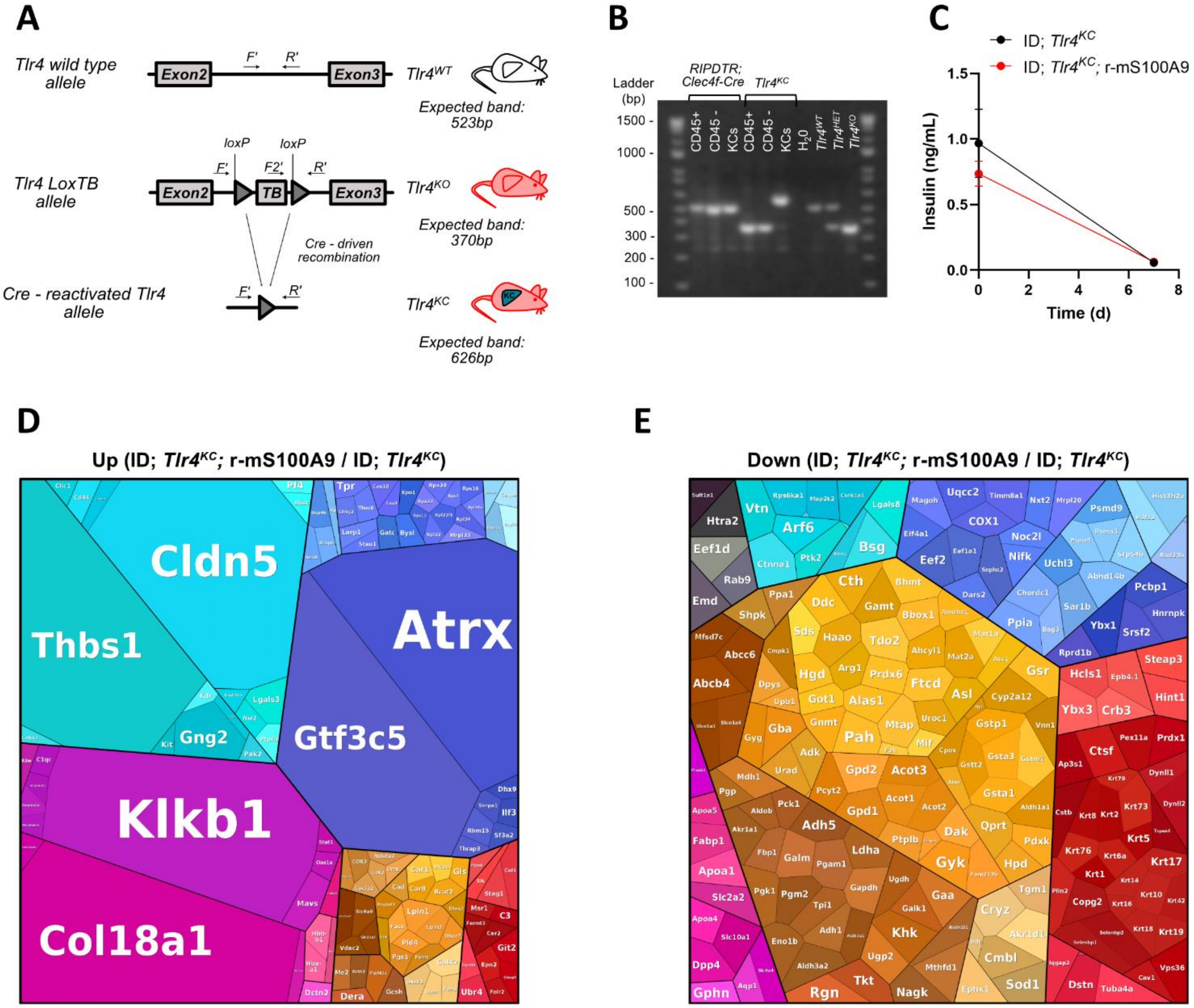


**Fig. S3. S100A9 modulates the KC proteome, promoting an anti-inflammatory and lipid-homeostatic signature in ID.**

**(A)** Schematic representation of the *Tlr4* wild type (top) and the *Tlr4*$^{LoxTB}$ allele before (middle) and after (bottom) Cre-recombinase-mediated deletion of the loxP-flanked transcriptional blocking (TB) sequences. F', F2' and R' are the primers used for genotyping, on the right, the expected amplicon sizes in PCR genotyping are indicated.

**(B)** PCR genotyping of the different cell fractions isolated through FACS sorting from *RIPDTR*; *Clec4f-Cre* and *Tlr4*$^{KC}$ mice.

**(C)** Plasma insulin was measured on day 0 and 7 in ID; *Tlr4*$^{KC}$ and ID; *Tlr4*$^{KC}$*;* r-mS100A9 mice as shown in Fig. 3A.

**(D-E)** Proteomaps illustrating in detail the changes in the KCs proteome upon r-mS100A9 treatment: treemap of proteins increased (fold change≥2) (**D**) or decreased (fold change≤0.5) (**E**) by r-mS100A9 treatment. Neighboring areas correspond to related functional groups and are marked by similar colors. The size of each polygon depicts the mass fraction of the respective proteins, reflecting protein abundances adjusted for their size.

Error bars represent SEM, statistical analyses were done using one-way or two-way ANOVA (Tukey's post- hoc test).

**Fig. S4.**

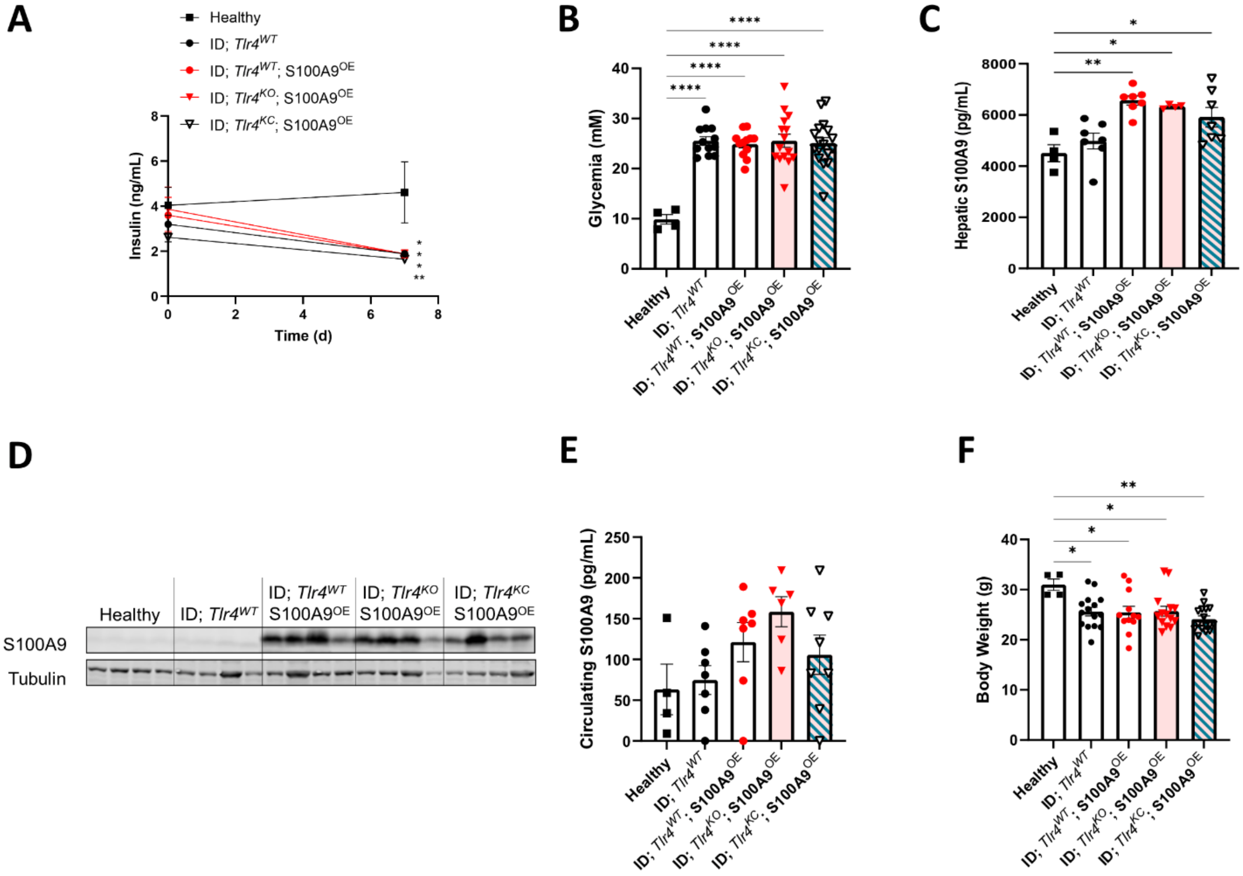


**Fig. S4. S100A9 rescues diabetic hyperketonemia and hypertriglyceridemia via TLR4 in KCs.**

**(A)** Plasma insulin levels of experimental mice as shown in Fig. 4A at day 0 and day 7.

**(B-C)** Glycemia (**B**) and hepatic S100A9 content (**C**) in the Healthy (n=4), ID; $Tlr4^{WT}$ (n=19), ID;$Tlr4^{WT}$;$S100A9^{OE}$ (n=15), ID;$Tlr4^{KO}$;$S100A9^{OE}$ (n=18), ID;$Tlr4^{KC}$;$S100A9^{OE}$ (n=16) groups measured on day 7 after 3h of fasting.

**(D)** Immunoblot of liver lysates for the S100A9 and Tubulin proteins of the indicated groups as shown in Fig. 4A on day 7.

**(E-F)** Circulating S100A9 (**E**) and body weight (**F**) measured on day 7 of indicated groups as shown in Fig. 4A.

Error bars represent SEM, statistical analyses were done using one-way or two-way ANOVA (Tukey's post- hoc test). Comparisons were made with Healthy controls unless otherwise specified on the graph. *$p \leq 0.05$, **$p \leq 0.01$, ***$p \leq 0.001$.

Fig. S5

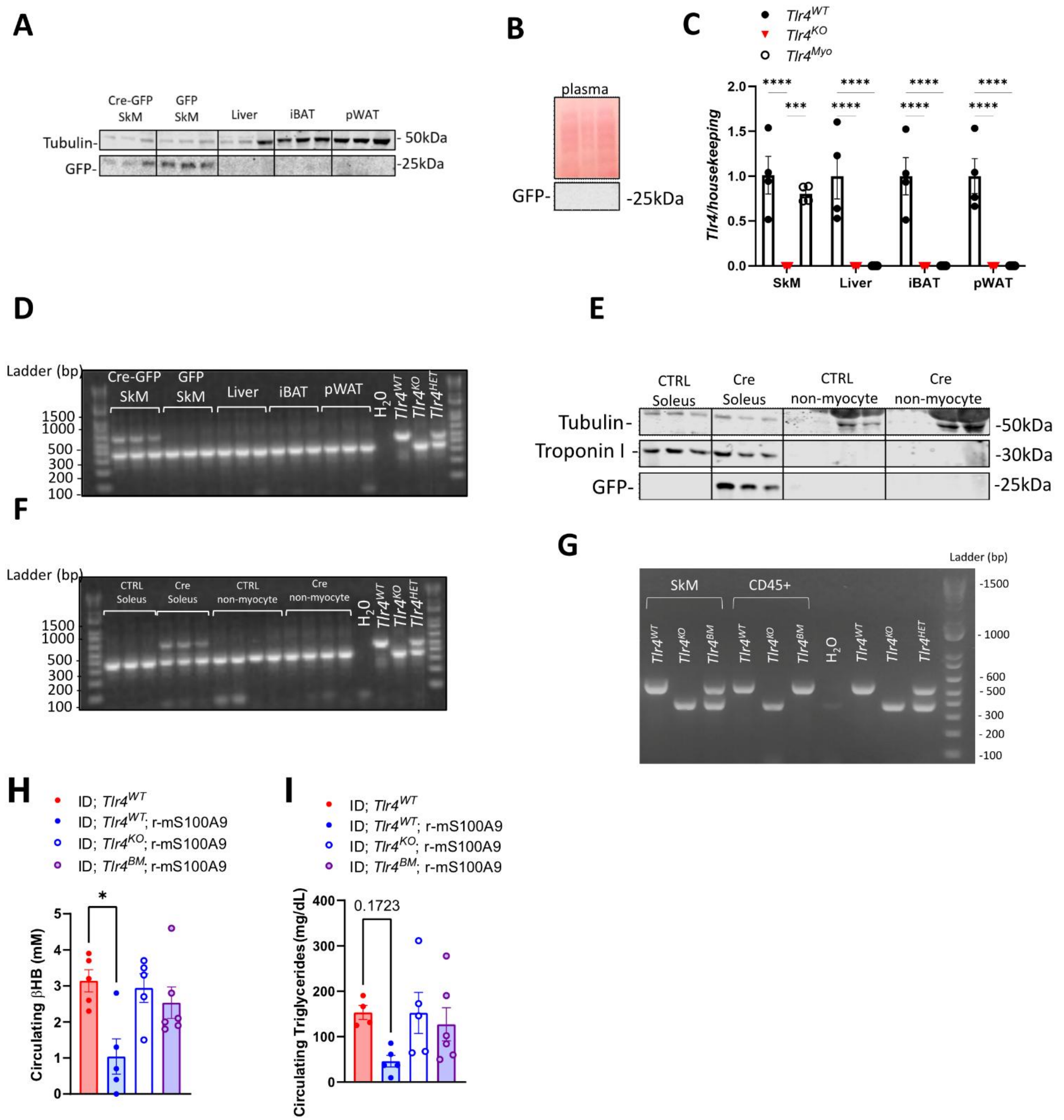


**Fig. S5. S100A9 exerts glucoregulatory effects *via* TLR4 on immune cells in SkM.** Hindlimbs of *Tlr4*$^{KO}$ mice were injected with 1x10^9 PFU of AAV expressing Cre-GFP (Cre-GFP SkM)or a GFP only control (GFP SkM) (n=3-4 per group) (*Tlr4*$^{Myo}$ mice). Blood and relevant tissues were isolated 7 days after injection.

**(A-B)** Immunoblots of GFP and Tubulin in the indicated tissues (**A**) and plasma (**B**).

**(C)** *Tlr4* mRNA content in the liver, SkM, iBAT, and pWAT of $Tlr4^{Myo}$ mice.

**(D)** PCR genotyping of *Tlr4* wild type and the $Tlr4^{LoxTB}$ allele before or after Cre-recombinase-mediated deletion of the loxP-flanked TB sequences (schematic representation of these alleles is shown in Fig. S3A) in indicated tissues from $Tlr4^{Myo}$ mice.

**(E)** Immunoblots of Tubulin, Troponin I and GFP in the Soleus and in the non-myocyte cell fraction from $Tlr4^{KO}$ and $Tlr4^{Myo}$ mice.

**(F)** PCR genotyping of *Tlr4* wild type and the $Tlr4^{LoxTB}$ allele before or after Cre-recombinase-mediated deletion of the loxP-flanked TB sequences (schematic representation of these alleles is shown in Fig. S3A) of whole muscle lysate and non-myocyte fractions isolated from whole muscle of injected leg (Cre Soleus) and non-injected control leg (CTRL Soleus) from $Tlr4^{Myo}$mice.

Bone marrow was isolated from $Tlr4^{WT}$ mice and injected into lethally irradiated $Tlr4^{KO}$ mice ($Tlr4^{BM}$). Two weeks after engraftment, blood, muscle fibres and immune cells were isolated from soleus muscle and subjected to PCR analysis.

**(G)** PCR genotyping of *Tlr4* wild type and the $Tlr4^{LoxTB}$ allele before or after Cre-recombinase-mediated deletion of the loxP-flanked TB sequences (schematic representation of these alleles is shown in Fig.. S3A) of the soleus (SkM) or non-myocyte cell fraction (CD45+) of the soleus from $Tlr4^{WT}$ and $Tlr4^{BM}$ mice.

**(I-J)** Circulating βHB (**I**) and triglycerides (**J**) content in ID; $Tlr4^{WT}$, ID; $Tlr4^{WT}$; r-mS100A9, ID; $Tlr4^{KO}$*;* r-mS100A9, and ID; $Tlr4^{BM}$*;* r-mS100A9 mice.

Error bars represent SEM, and statistical analyses were done using one-way ANOVA(Tukey's post hoc test). * indicates comparison to $Tlr4^{KO}$ allele. *$P < 0.05$, **$P < 0.01$.

**Table S1.**
Excel file containing the transcription factor enrichment data, related to Fig. 2.
